# Nanoscopy on the Chea(i)p

**DOI:** 10.1101/2020.09.04.283085

**Authors:** Benedict Diederich, Øystein Helle, Patrick Then, Pablo Carravilla, Kay Oliver Schink, Franziska Hornung, Stefanie Deinhardt-Emmer, Christian Eggeling, Balpreet Singh Ahluwalia, Rainer Heintzmann

## Abstract

Super-resolution microscopy allows for stunning images with a resolution well beyond the optical diffraction limit, but the imaging techniques are demanding in terms of instrumentation and software. Using scientific-grade cameras, solid-state lasers and top-shelf microscopy objective lenses drives the price and complexity of the system, limiting its use to well-funded institutions. However, by harnessing recent developments in CMOS image sensor technology and low-cost illumination strategies, super-resolution microscopy can be made available to the mass-markets for a fraction of the price. Here, we present a 3D printed, self-contained super-resolution microscope with a price tag below 1000 $ including the objective and a cellphone. The system relies on a cellphone to both acquire and process images as well as control the hardware, and a photonic-chip enabled illumination. The system exhibits 100*nm* optical resolution using single-molecule localization microscopy and can provide live super-resolution imaging using light intensity fluctuation methods. Furthermore, due to its compactness, we demonstrate its potential use inside bench-top incubators and high biological safety level environments imaging SARS-CoV-2 viroids. By the development of low-cost instrumentation and by sharing the designs and manuals, the stage for democratizing super-resolution imaging is set.

## 1 Introduction

Driven by the desire to learn more about the behaviour of biological cells, microscopy has entered a race to resolve ever smaller structures. This facilitates the study of biological processes on a sub-cellular level, e.g. helping to understand the mechanisms of the human immune system [1, 2, 3] and the infection mechanisms of pathogens like viruses or bacteria [4, 5]. Optical, fluorescence-based methods have been established as a popular part in the toolbox of modern biological and medical research. As an important advantage, they can combine spatial information (i.e. fluorescence microscopy) with the capacity for sensitive and highly specific detection of antigens. The last decade, in particular, has seen the rise of optical super-resolution imaging (SRI), i.e. optical microscopy well beyond the resolution limit of *λ/*2 as described famously by Ernst Abbe [6]. Optical super-resolution (SR) can be classified into deterministic and stochastic methods. The former includes techniques based on point-scanning, e.g. stimulated emission depletion (STED) microscopy [7] or structured illumination microscopy (SIM) [8]. The latter comprise techniques such as Photoactivated Localization Microscopy (PALM) [9] or (*direct*) Stochastic Optical Reconstruction Microscopy (*d* STORM) [10, 11]. The SRI methods mentioned here seem to follow an inverse relationship between optical resolution and complexity of the optical setup. High costs for modern high-end microscopy and camera hardware as well as their often necessary stationary operation additionally often limit the access considerably.

Especially biologically difficult situations show how important SRI is for the investigation of particles such as the novel SARS-CoV-2 coronavirus with a size far below the optical resolution limit. Broad access to powerful nanoscopic imaging platforms in clinics and research institutions would have far-reaching implications for mankind by facilitating the detection and study of this pathogen. However, due to the cost, size and complexity, this is not possible with the current state of the art.

Several attempts already showed how to significantly reduce high costs and resulting exclusivity and aimed to provide greater accessibility of SRI [12, 13, 14]. Substituting expensive electron multiplied charged coupled device (emCCD) cameras by industry-grade [15] or even cellphone cameras [16], as well as relying on entertainment lasers [13] and compensating the lack of imaging quality with tailored image processing [17, 18, 19, 20, 21, 19, 22] contributed greatly to the idea of widely available high-resolution imaging platforms on a budget. Open-source 3D-printed approaches like the Openflexure[23], Attobright [24], Micube [25] or the 3D-printed optical toolbox UC2 [26] can further provide flexible and open hardware solutions in order to support software-based solutions. Alternatively, (mass-produced) photonic integrated circuits (PICs) enable the creation of compact, low-cost and sensitive optical setups. On-chip *d* STORM has exhibited SRI over a large field of views (FOV) [27] and on-chip total internal reflection fluorescence (TIRF) SIM has demonstrated capabilities to provide 2.4× full width at half maximum (FWHM) reduction of the point spread function (PSF) in linear SIM [28].

To unleash the full potential of photonic chip-based nanoscopy to support the ongoing SARS-CoV-2 pandemic, we combine chip-based illumination and cellphone-based imaging into a compact and affordable device “*cell*STORM II” which delivers images down to 100*nm* and can operate autonomously over long periods of time. To accomplish that, we employ the mobile phone not only as a camera but for simultaneous on-device data evaluation by implementing neural network-based (NN) image processing of the camera stream. A key enabler here is the development of an affordable optomechanical architecture based on optical pick-up units (OPU) of optical disk players (e.g. Bluray) to efficiently couple light into the photonic chip. We have further developed a customized application (APP), STORMimager, that can run on Android cellphones and controls the entire pipeline of the proposed on-chip nanoscopy.

Three use cases demonstrate the impact of this ≤ 1000 $ SRI device in areas where highly sensitive images are required but access to the necessary tools is often very limited. A benchmark of the optical resolution of intensity fluctuating SR and SML methods (e.g *d* STORM) benchmarks the optical resolution. The ability to perform long-term imaging of living cells within cell incubators and the possibility to use the device in high biological safety level environments such as BSL3 laboratories enables the investigation of dangerous pathogens, as demonstrated on immunostained SARS-CoV-2 virus particles *in-vitro* and a colocalization of fixed pseudotyped lentiviruses (i.e. mCherry and immunostained) to determine the binding specificity of the antibody.

To enable wider penetration of this powerful imaging methodology, all sources, schematics, blueprints, tutorials for simple replication and the software are publicly available in our Github repository (https://beniroquai.github.io/stormocheap/).

## 2 Results

### 2.1 Autonomous, Compact and Low-Cost Super-Resolution Imaging Device

Fluorescence imaging and SMLM, in particular, place high demands on both proper sample preparation and the signal to noise ratio (SNR) to obtain high-resolution images. Especially, SMLM with *d* STORM requires a high excitation intensity (i.e. *>* 1*kW/cm*^2^) to produce the desired blinking behaviour that allows the localization of single emitters below the diffraction limit [11]. To achieve this within the evanescent field of planar waveguides, a high coupling efficiency is beneficial. So far this has been achieved by using high-end lasers and focusing optics in combination with a precise and costly piezoelectric alignment stage [27]. We noticed that many consumer electronics with electro-optical components such as cellphone cameras or DVD-players contain a miniaturized version of similar arrangements in the form of adaptive optical elements. Most optical image stabilization (OIS) system in cellphones [29] or optical pickup units (OPU) with auto-focusing systems inside optical disk players are realized by moving a lens inside a voice coil motor (VCM) [30, 31]. This allows rapid (bandwidth 20 *kHz*) and precise movements (nanometer resolution, 1 *mum/mV*) to compensate motion artefacts and to ensure constant optical readout using closed-feedback loops [32]. We build upon this feature and choose a widely available, thus reproducible OPU (KES400A, Sony, Japan) from a Bluray player of a scrapped Playstation 3 (Sony, Japan, 1-5 $ for the spare OPU) as an alternative to common nano-positioning fiber coupling systems (e.g. Nanomax, Thorlabs, USA, 2500 $).

To have a compact, but also low-cost optical detection path using off-the-shelf components, we decided to rely on a compound microscope design equipped with an autofocusing mechanism (see Methods 3.1). A monolithically printed part defines all-optical relationship (red part in Fig. 1b) and enables fast reproduction of the setup.

**Fig 1.**
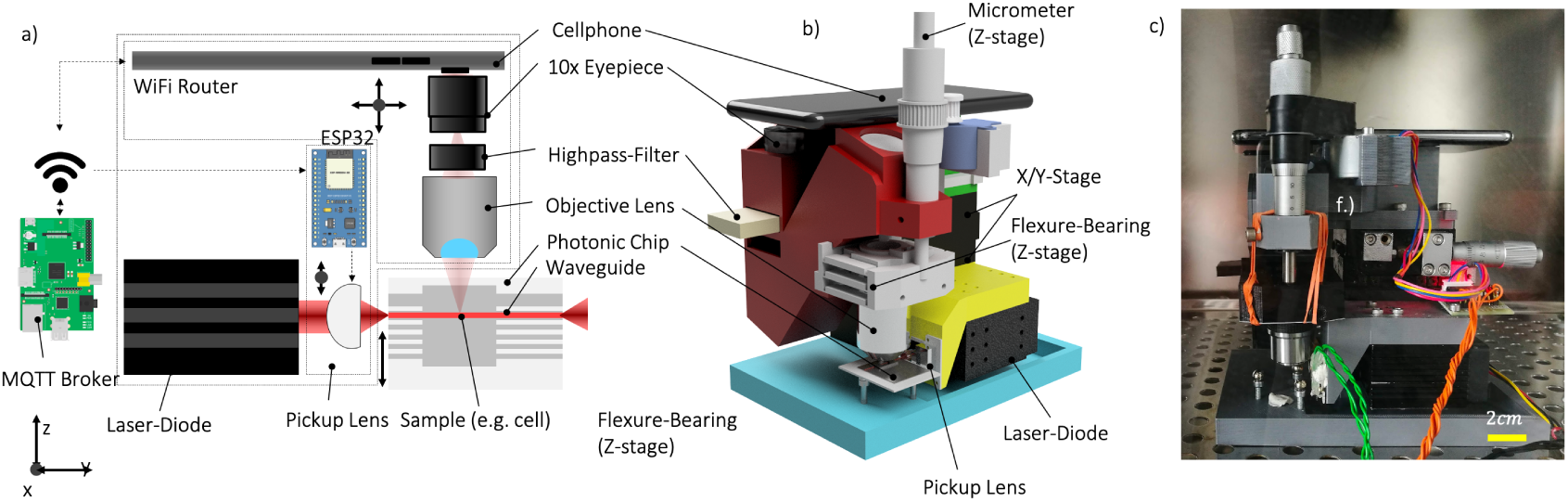
Portable SMLM Setup. a) The two XY-stages realize coarse, the voice-coil motor (VCM) support fine alignment of the focused laser relative to the stationary waveguide chip as well as moving the optical module to select the field of view (FOV). The sample is observed in TIRF from above, b) the CAD drawing shows the device where the cellphone controls all actuators and performs imaging. A motor-driven flexure bearing-based focusing mechanism allows for precise focusing, c) the small footprint of the assembled microscope enables convenient placement inside biological incubators or laminar flow hoods

To achieve the highest possible coupling efficiency, precise placement of the lens relative to the rigidly mounted multimode waveguide (WG) is required. Rough coupling with the XY-Stages (No Name, 80 $, China) is followed by fine coupling with the current-controlled voice coil-mounted coupling lens of the OPU in the sub-micrometer range. This, as well as the focus-position and laser intensity, can be controlled manually or automatically by an auto-adjusting algorithm using a customized Android APP (available at [33], Methods 3.2).

While the transverse lens movement (along z-axis in Fig. 1a) was mostly sufficient to achieve good coupling, rapid axial motion (along y-axis in Fig. 1a) enables both the destruction of the static mode field pattern produced by the mode-selective behavior of the WGs [28, 34] at frequencies higher than the pixel integration time (e.g. ≥ 500 *Hz*) or the realization of fluctuating intensity-based super-resolution methods such as SOFI [35], SRRF [18] or ESI [21] at lower-frequencies (e.g. ≤ 10 *Hz*), where each frame sees a different emitted signal. We build upon *cell*STORM as previously described in [16] and take advantage of the phone’s full potential in terms of its camera, computational resources and ability to communicate wirelessly. The possibility to control hardware components based on the results of the on-device image processing enables the creation of direct feedback loops for automatic WG coupling (see Supp. Notes S1.3) and autofocusing (see Supp. Notes S1.2). This compensates for any drifts and thus allows for long-term unattended operation, such as imaging inside incubator. By taking a cellphone (Huawei P20 Pro, China, 300 $) equipped with a monochromatic back-illuminated CMOS sensor (Sony IMX600 mono, 5120 × 3840 pixels^2^, 1.01*µm*, Japan, for effective pixel sizes see Supp. Notes S1.8) we take advantage of higher photon budget compared to RGB-sensors (using Bayer patterns).

Further, we demonstrate how the immense computational power of the cellphone ensures an improvement in image quality without transferring data to external computers. Similar to the methods SOFI [36, 34], SRRF [18] or ESI[21] we want to take advantage of mobile phone optimized image processing libraries like the machine learning framework “Tensorflow Lite [37, 38] to calculate instantaneous super-resolution images from intensity fluctuations caused by a variation of the waveguide mode pattern. We aim to preserve the temporal relationship of acquired frames by designing the model using convolutional Long Short Term Memory (*c*LSTM, [39]) layers. This architecture (Fig. 2a) initially used for applications such as weather-forecasting [40] and future frame prediction [41], is ideal for 2D time-series, where information along *t* is correlated. The neural network takes a series of TIRF images (e.g. 20 frames) at varying excitation pattern (e.g. randomly oscillating OPU) and extracts spatial features like edges in each time-step. The hidden state of the recurrent network preserves temporal relations such as the correlated signal of an illuminated fluorophore which is assumed to be independent of noise [42]. The extracted features are merged by an adjacent convolutional layer before they get up-sampled using a sub-pixel convolutional-layer [43]. The result is a two-fold “super-resolved” image overlaid on the video stream of the microscope screen (see Supp. Video S1.2). The network’s architecture is a compromise between mathematical operands running on the cellphone and performance in terms of processing speed and resolution improvement.

**Fig 2.**
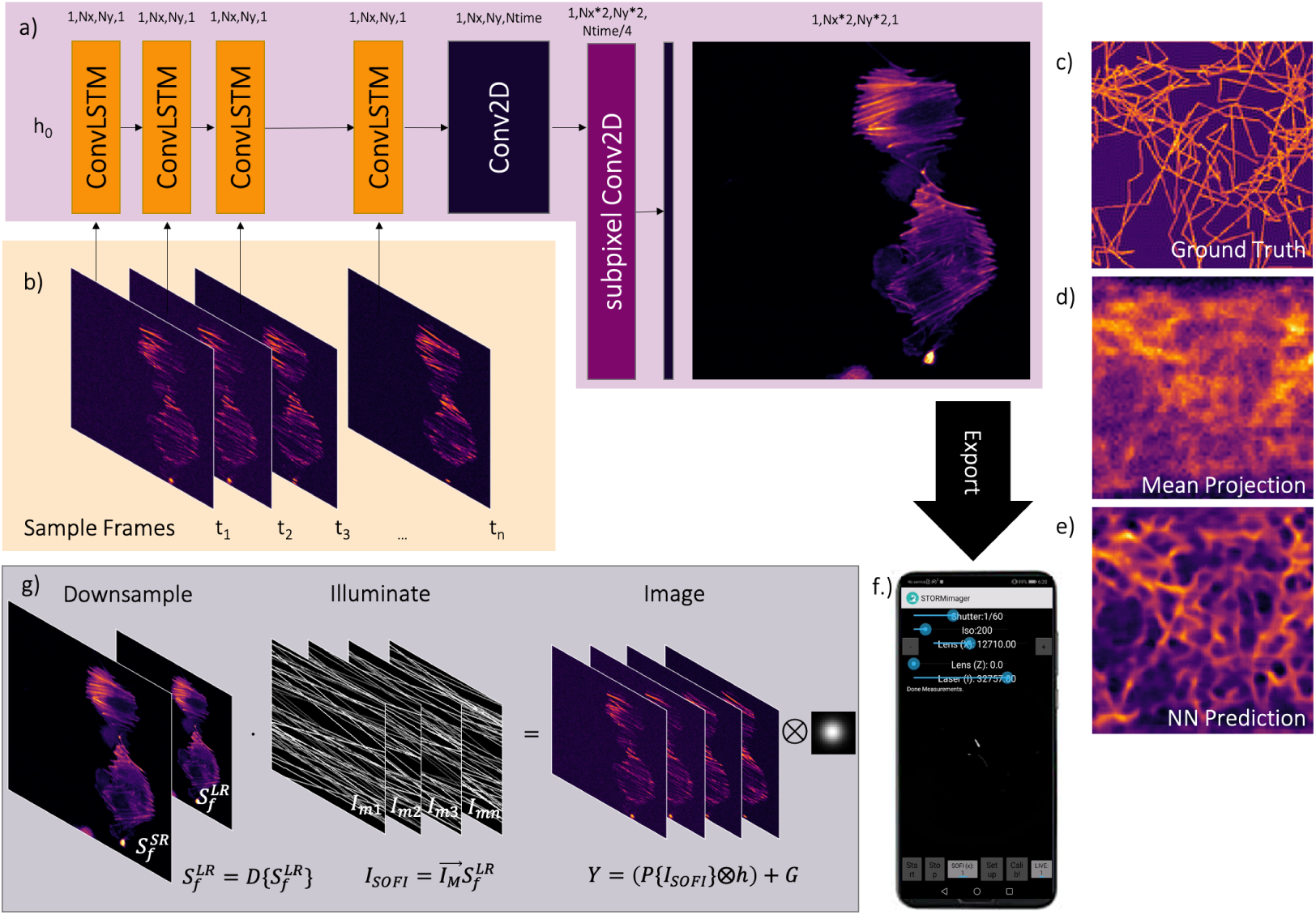
Neural Network Architecture. a) the sequential convolutional long short term memory (cLSTM) neural network receives a series of fluorescent images b) with varying excitation pattern and predicts a “super-resolved” representation e) which outperforms variance and mean projections d). The pre-trained model can be ported and deployed on the cellphone f) to yield an instant super-resolution image which allows interrupting failed experiments quickly. The training dataset is generated by simulating the process of imaging at fluctuating excitation intensity, where a high resolution fluorophore distribution 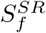 is first downsampled, before is illuminated by an artificially created temporal varying excitation pattern 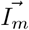 and imaged by the microscope model *H*{·} (see Methods 3.2). A time-series of *n* = 30 frames of the ground truth distribution of fluorophores c) using the model in g) lead to the prediction in e). It outperforms the mean projection d), where many structural details are lost.

Similar to work in [44] we create a training dataset which contains realistic TIRF images and their SR equivalents. The model *O*{·} visualized in Fig. 2 g) imitates the acquisition process of a time-series at varying excitation pattern by synthetically illuminating a spatially varying fluorophore distribution *S*_*f*_ with a random line-like excitation pattern 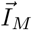. The temporally changing excitation pattern creates a stack which is further processed by a realistic microscope model, where noise and low-pass filtering is applied (see Methods 3.2). Combined with the automatic coupling mechanism, this is a step towards an “intelligent” microscope placing less burden on the user as additional parameters for optimal control of the instrument as well as data read-out are handled automatically (see Section 2.2).

### 2.2 Improving Cellphone-based SRI using RAW Imaging Data in SRRF and *d* STORM experiments

In our previous study *cell*STORM we identify unpredictable post-processing of image data by the post-processing performed by the smartphone firmware leading to a reduced optical resolution or artefacts in the final reconstructed image [45, 16] as a serious problem of smart phone-based super-resolution microscopy. To give an example, low signal blinking events may accidentally be removed by the denoiser before the adjacent video compression alters the data by finding a sparse representation in the spatial and temporal domain. In case of the H264 codec an integer transform (*IT*, [46]) reduces the information bandwidth in each frame before information between consecutive frames is reduced by interpolating between several intra or key- (I), predictive- (P) and bidirectional- (B) coded frames. In case a sampled PSF from a single blinking event is located close to the edge of an *IT* block, the H264 cannot reproduce the rotational-symmetric shape (e.g. Gaussian) properly. The result is a discontinuity to neighboring blocks, which results in a destruction of localization accuracy and the production of the grid pattern in the reconstruction (e.g. ThunderSTORM, SRRF, Supp. Fig. S1.1). The temporal interpolation results in unpredictable and unrecoverable shifting of events between frames, therefore representing an additional error source especially visible in low-SNR situations (e.g. *d* STORM measurements). For a qualitative comparison between compressed and non-compressed data see Supp. Image S1.2).

To solve the associated problem of introduced reconstruction artifacts, we enabled RAW image acquisition at video-rates (e.g. 5-10 frames per second, FPS) at 200 × 200 pixels (see Sect. 3.3). In Fig. 3 we compare compressed and uncompressed measurements with static widefield (WF) and fluctuating illumination which are further processed using SRRF [18] on a GPU-equipped desktop computer and the proposed NN directly running on the cellphone. Additionally, we perform a benchmark-test using *E. coli* bacteria stained with mCLING-ATTO647 as a reference sample to compare between the cellphone-based device (100×, *NA* = 1.25 oil, China, 80 $) and a standard research microscope (Zeiss Axiovert TV, Germany) equipped with a TIRF lens (Zeiss *α* Plan-Apochromat 100× */*1.46).

**Fig 3.**
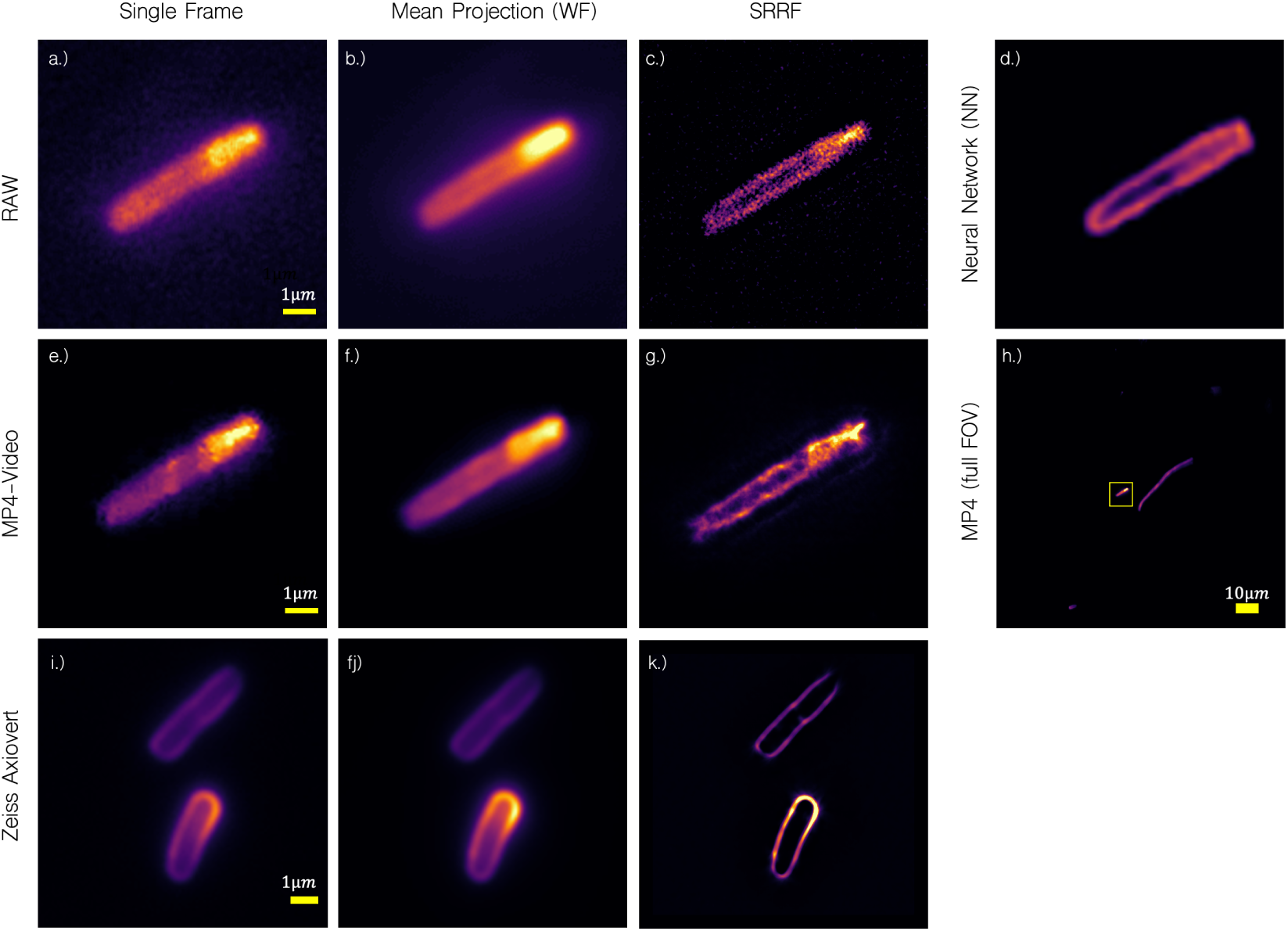
SR Reconstruction of Compressed and Raw Measurements and Fluctuating Intensity Illumination. a) Image series in “RAW” mode exhibit represent the raw unaltered value of the camera sensor and excludes post-processing such as denoising or compression which is e.g. visible as block-like artefacts in the MP4 video frame e) also resulting in grid-like artefacts in the SRRF reconstruction g), not visible in the reconstruction of RAW data c). Processing the data with SRRF (RAW: c); MP4: g)), as well as with the experimental NN running at 1 *fps* directly on the device d) successfully recover the membrane of the *E. coli* bacteria. Nevertheless, video-series imaging has the clear advantage of an increased FOV depicted in h) and Supp. Video S1.4, where the yellow box is the cropped area for e)-g). A comparison of the same sample using a research-grade microscope (Zeiss Axiovert TV, 100×, *NA* = 1.46) demonstrates, that the smartphone-based microscope can compete with the inverted TIRF imaging device more than one order of magnitude more expensive.

The transversally oscillating OPU in case of the *cell*STORM II and a slight variation of the excitation beam inside the back-focal plane (BFP) in case of the Zeiss microscope resulted in sufficiently random fluctuation of the excitation intensity to process the data with SRRF. The cameras exposure time (e.g. *t*_*exp*_ = 1*/*200 *s*) and gain (e.g *ISO* = 1000) were tuned to achieve a reasonable high frame rate (e.g. 30 *fps*) at a high SNR. Figure 3 shows a comparison of a single frame of a time-series, their mean-projection and the super-resolved result using SRRF of RAW and compressed measurements.

Even a single frame from a compressed video-stream (Fig. 3e) exhibits several alterations compared to its RAW frame equivalent (Fig. 3a). This results in grid-like artefacts in the SRRF reconstruction (see also Suppl. Fig. S1.1), causing an “oscillating” bacteria membrane (Fig. 3g), which appears as a straight line in the reconstruction of 200 RAW frames (Fig. 3c) recorded with the Zeiss Axiovert (Fig. 3k) and as known from the literature (e.g.[47]). This is also highlighted in the measurement using the Zeiss Axiovert shown in Fig. 3i) which offers the same amount of details compared to the low-cost cellphone-based device, also manifested in the Fourier ring correlation (FRC) [48] determined resolution of SRRF reconstructions which is *d*_*Zeiss*_ = 158*nm* (Fig. 3k) for the Zeiss microscope and *d*_*RAW*_ = 122*nm* (Fig. 3c) and *d*_*MP*4_ = 320*nm* (Fig. 3g) for cellphone respectively. As previously mentioned, SRRF adds a high-frequency pattern to the reconstructed video data (see Supp. Figure S1.1) which is falsely detected as the object structure by the FRC and gives the impression of a higher resolution. Measuring the thickness (FWHM) of the bacteria membrane leads to a resolution of *d*_*Zeiss*_ ≈ 160 *nm, d*_*MP*4_ ≈ 210*nm* and *d*_*RAW*_ ≈ 250 *nm*. Although we are not directly aiming for super-resolution imaging with the parameter-free neural network, we were able to successfully recover membrane in measurements with *E. coli* bacteria (see screenshot in Fig. 3d) which results in a resolution of *d*_*NN*_ ≈ 240*nm* (line plot). The software running in the smartphone APP STORMimager [33] creates a real-time screen overlay (i.e. 1 *fps* at 200 × 200 pixel^2^, Supp. Video S1.2) and opens the possibility to check experimental parameters directly in the lab without the use of external software like Fiji [49].

Although the compressed video stream (e.g. 2000 pixels^2^) leads to reconstruction artefacts, the 100-fold increase of the FOV (Fig. 3h) compared to RAW measurements (e.g. 200 pixels^2^) at a sufficiently high frame rate underlines the advantage of compact nanoscopic imaging of large FOVs on photonic chips. This is especially true for *d* STORM experiments, where this setup grants access to high optical resolution at large FOVs. Therefore, we tested the proposed setup in a set of *d* STORM experiments, where we labelled the actin network of different cells such as HeLa and U2OS with Alexa Fluor 647 Phalloidin (Thermofisher, MA, US) captured with a 60×, *NA*=0.85 (Fig. 4a-c) and 100×, *NA*=1.25 oil (Fig. 4d) objective (both No Name, China, 65 $). Using Oxyrase/*β*-Mercapto Ethylamine (OxEA) as a *d* STORM buffer [50] and optimizing the coupling intensity, we achieved good switching efficiency between dark and bright states, while observing relatively low photobleaching during the experiment (See Suppl. Video S1.3). Video acquisition during experiments was performed with the APP FreedCam [51] (e.g. *t*_*exp*_ = 1*/*20 *s, ISO* = 2000) on an optical table to reduce low-frequency vibration. The lateral focus drift of up to 5 *µm* (see. Graph in Supp. Notes S1.2) was compensated using image post-processing inside the localization software ThunderSTORM [52] or the Nano-J suite [53] in Fiji [49]. A second MQTT-enabled smartphone acting as a remote control enables the coupling efficiency and focus position to be adjusted without touching the imaging device.

**Fig 4.**
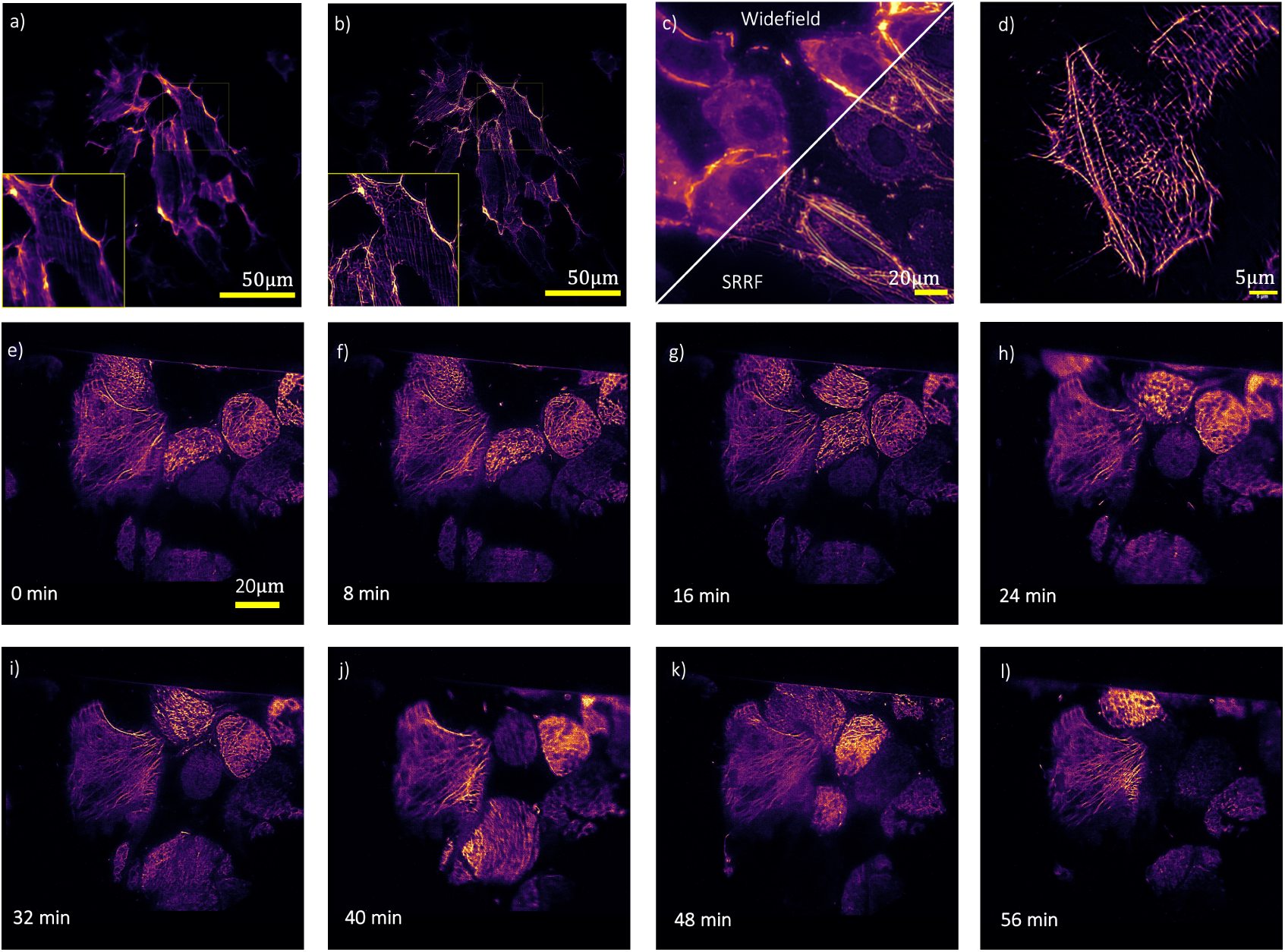
Versatile Tool for Biological Imaging. a) Is the widefield equivalent as the temporal sum of a *d* STORM experiment of AF647 Phalloidin labelled HeLa reconstructed using ThunderSTORM in b), where the Actin filament network is clearly recovered over a very large FOV (≥ 200 *µm*). Localization accuracy is degraded by field curvature at the edges of the FOV. c) Another comparison of widefield and its SRRF reconstruction of blinking U2OS cells. d) An additional resolution improvement is achieved by using high *NA* immersion objective lenses (i.e. 100×, *NA* = 1.25), which produces ≤ 100*nm* optical resolution. d-l) SRRF reconstructions of a long-term (17 *h*) measurement of SiR-labelled HeLa cells *in-vitro* at fluctuating excitation intensity (i.e. oscillating coupling lens). The unsupervised experiment was performed at room temperature to study cell movement. In each time-stamp (*t* = 4 *min*) the Actin filament network is clearly recovered and shows strong cell dynamics in the first few minutes.

With this method, we achieved a resolution of up to 127*nm* measured with FRC (Fig. 4 d) and *<* 100*nm* using line plot. Nevertheless, the additional gain in resolution in *d* STORM measurements is relatively small compared to the previously measured improvement from fluctuation-based measurements which also enables live-cell imaging.

### 2.3 Versatile Tool for Multimodal *in-vitro* TIRF Imaging in Biological Labs

Bio-containment regulations constitute one of the main limitations to the introduction of advanced microscopy methods into environments where infectious pathogens are investigated. Biosecurity precautions discourage the use of expensive commercial microscopes in these laboratories, e. g. due to the restriction to remove live samples from laminar flow hoods or the necessity to thoroughly disinfect any material before using it in an environment of lower containment level. In particular, disinfection procedures can potentially damage delicate optical components. The small footprint and its autonomous operational mode of the self-contained microscope enables its use in these environments, which involves the use in bench-top incubators and in places where laboratory equipment is difficult to obtain.

Since important parts of our microscope design are 3D-printed it lacks long-term stability, especially when temperature fluctuation leads to a deformation of the thermoplastics (e.g. Polylacatid acid, PLA). This is especially visible during long-term experiments in biological cell incubators, where heat and humidity (*T* = 37°*C, CO*_2_ = 5%, Humidity=100 %) lead to large temperature-dependent drift (lateral and axial). The aforementioned autofocus (see Supp. S1.2) inside the customized APP “STORMimager” [26] in combination with a mechanical design which minimizes drift can successfully compensate for that and enables live-cell imaging. This is demonstrated by placing the whole setup inside an incubator (Thermofisher Heracell, MA, USA) and observing Mitotracker Orange (Thermofisher, MA, USA) labelled U2OS cells for 17 *h*. Unfortunately, the cells did not adhere well to the surface of the chip and therefore did not yield useful results. Additionally, we performed an overnight measurement of silica rhodamine-actin (SiR-actin) [54] stained HeLa cells at room temperature to observe apoptosis (see Supp. Video S1.1). Periodic measurements (Fig. 4e-l) at fluctuating intensity give access to a higher optical resolution and recovers the actin filament. All electronics, including the mobile phone, can withstand the environmental conditions inside incubators that are outside of their intended use for multiple days.

We then set out to demonstrate the use of the *cell*STORM II system for advanced biological research questions. Antibody-body based microscopic detection of pathogens or infected cells remains an important diagnostic tool [55]. In addition, there is a need for high resolution microscopy in many labs working with viruses and other pathogens. However, installation of advanced microscopes in high security laboratories is often challenging and requires extensive adaptions of the microscope to allow safe work with pathogens. In contrast, the *cell*STORM II system is small enough that it can be directly installed in biosafety cabinets and thus can be used in high safety laboratory settings. Alternatively, wave-guide-based high resolution microscopes could form the basis for fast and cheap diagnostic instruments, which could serve as point-of-care (PoC) diagnostic tools, especially in countries with poor laboratory infrastructure. We therefore tested if the system can directly detect viral particles by immunofluorescence. As a model pathogen, we chose the newly discovered SARS-CoV-2 coronavirus. To test if the *cell*STORM II system can be used in a high security biological laboratory, we installed one system in a biosafety level 3 (BSL3) laboratory at the University clinic Jena (UKJ, Germany) and tested if we can detect viral particles by immunofluorescence. To this end, we immobilized active SARS-CoV-2 virus particles on the surface of the wave guide and stained the particles with anti-SARS-CoV-2 antibodies. Using this setup, we were able to detect viral particles and – using *d* STORM – were able resolve them as spherical particles.

We noted that the number of observed particles by microscopy - as determined by counting antibody foci - was significantly higher than the number of active SARS-CoV-2 infectious particles determined by plaque assays. While this may be explained by the fact that not all formed viral particles are functional and infectious, we performed additional tests to exclude the unspecific binding of the antibody to the waveguide surface. To this end, we generated virus-like particles (VLPs) which were pseudotyped with either the SARS-CoV-2 Spike protein or the native HIV-1 Env protein. The SARS-CoV-2 Spike protein was a mixture of GFP-tagged and non-tagged S. In addition, we used MyrPalm-mCherry (SARS-CoV-2 S-containing VLPs) or GFP-labelled Vpe (HIV Env VLPs) to label individual particles. This allowed us to unambiguously identify viral particles and test if the antibody bound specific to them or exhibited unspecific binding to the wave guide surface.

We then applied the same immunohistological protocol as for the active SARS-CoV-2 viroids and used the two-color extension of the setup (see Supp. S1.7) with *λ*_*mCherry*_ = 532*nm* and *λ*_*AF*647_ = 635*/*637*nm* to detect SARS-CoV-2-pesudotyped VLPs, and the same setup with an additional laser *λ*_*eGFP*_ = 488*nm* to detect HIV Env pseudotyped VLPs. In the case of the SARS-CoV-2 pseudotyped VLPs, we observed multiple antibody-positive foci. We also observed a number of non-colocalizing signals, which can be explained that integration of either Spike protein and MyrPalm-mCherry is a stochastic event, and not every VLP formed necessarily contains both of them. In contrast, when we labelled GFP-labelled HIV Env-pseudotyped VLPs with the SARS-CoV-2 antibody, we did not observe any colocalization, indicating that the observed antibody labelling is specific. To give a quantitative measure about the binding specificity, we quantified the antibody intensity in regions containing VLPs (mCherry/EGFP positive) and non-viral(background, mCherry/EGFP negative) regions as described in [58]. Figure 5g). This quantification showed that the antibody signal around SARS-CoV-2 VLPs was on average 8.7× higher than in background regions,whereas we observed no enrichment (0.9 ×) in the case of HIV-1-pseudotyped VLPs. This result highlights the potential of *cell*STORM II for viral research.

**Fig 5.**
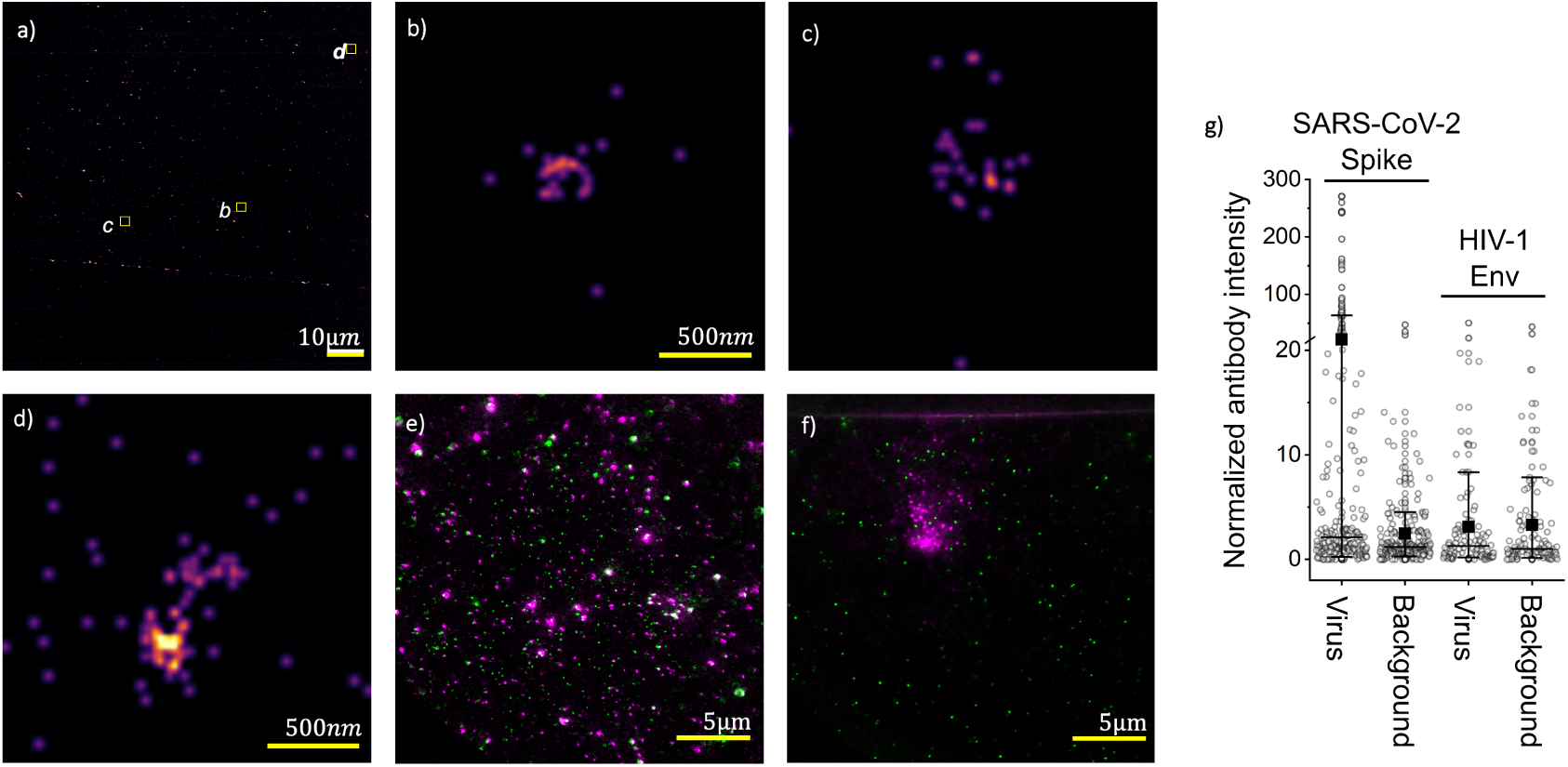
Resolving of Pathogens SARS-CoV-2. The setup can be placed in high biosafety areas such as BSL3 labs, where we resolved a) active SARS-CoV-2 virus particles using *d* STORM. b)-d) show zoomed-in ROIs where the appearance of the particles follow the expected shape (e.g. spherical) and size (≈ 150 *nm*). To justify the binding specificity of the SARS-CoV-2 *α*-spike antibody, we performed a series of colocalization experiments on the *cell*STORM II device using its dual-colour mode. e) Each green spot represent a signal from mCherry expressing pseudo-typed Lenti SARS-CoV-2 viruses expressing the spike protein, where the purple spot comes from the *α*-spike antibody labelled with anti-mouse AF647. f) The control of pseudo-typed HIV particles expressing GFP show much lower signal from the same *α* spike antibody. g) Intensity quantification of images e) and f) in viral (mCherry positive for SARS-CoV-2 VLPs and EGFP positive for HIV-1 VLPs) and non-viral (background, mCherry/EGFP negative) ROIs, both 5 pixels in diameter. Viral ROIs were identified by a local intensity maxima detection algorithm. Then, the intensity of the Alexa Fluor 647 channel was quantified in these ROIs. Each point corresponds to the intensity normalized to the background median value for each virus or background ROI, black squares represent the mean, horizontal lines are the median and whiskers are the 10^th^ and 90^th^ percentile.

As mentioned previously, SMLM is capable of resolving the spherical corona viruses (approx. ∅ = 120 − 160 *nm*, [57]) in comparison to wide-field fluorescence microscopy in order to provide an additional parameter for evaluation. The zoomed-in *d* STORM reconstructions in Fig. 5 b-d) show (hollow) spheres at a size corresponding to the virus particle between (120 − 180*nm* in diameter). Due to heavy air ventilation inside the laboratory and low signal after about 5 *min* of acquisition time due to photobleaching, the reconstruction is based on ≈ 3000 frames only. Per-frame drift correction was performed with the NanoJ toolbox [53], where we used strong inelastic scattering at the edge of the waveguide as fiducial markers (straight line in 5 a). This, together with the low penetration depth of the evanescent field of the WGs of about 150*nm* increases the sensitivity of the measurement and increases the probability that the detected particles are actually SARS-CoV-2 viroids.

## Discussion

With *cell*STORM II we are pursuing ideas initially proposed in [28] and [16] to create a low-cost and portable cellphone-based SMLM device. We show that recent rapid-prototyping methods in combination with integrated photonic waveguide chips and open-source hard- and software enable the creation of SRI imaging devices for applications, where common microscopes are hardly available. Imaging modalities such as *d* STORM in fixed cells at high optical sectioning (i.e. TIRF) and large field-of-views and even live-cell imaging using fluctuation-based approaches open up new opportunities for biological research. The small footprint and overall low price allow for quick replication and transportation to different locations at remote places or with high-safety restrictions (e.g. BSL3 laboratory). Components which are broken during experiments (e.g. due to material fatigue) can easily be replaced or upgraded, thus ensuring a long lifetime of the device and the potential to adapt different techniques, exemplarily shown with the multicolour imaging or automatic coupling routine. With this, *cell*STORM II is promoting reproducible science for many people leading to further democratization of SR imaging tools. The fact that rapid and frequent changes in the software and hardware components of mobile phones can cause a wide variation in image quality, puts the use of recently introduced edge computing devices (e.g. Nvidia Jetson Nano, USA, Raspberry Pi, UK) at the centre of future developments. These computationally powerful devices in combination with highly sensitive image sensors from the field of embedded vision are available in large quantities on a long-term basis. This way they always offer the same quality and thus a high degree of reproducibility, autonomy and quality at a very reasonable price.

The setup design is the result of an iterative optimization between available, yet inexpensive components and high optical quality. The overall design of the microscope highlights several problems which can potentially be solved with a higher budget. For the compound microscope-based optical design for instance most “budget” objective lenses, in general, perform well around their centre FOV, but show increased aberration and field distortion at the edges. This and a tilted image plane which origins in a possible angle between the WG chip and the objective lens leads to an according decrease in reconstruction accuracy. This, in turn, compromises the advantage of a large FOV in on-chip imaging. Additionally, we found that different coupling lenses from CD or DVD drives may show slightly better coupling performance due to a better matching centre wavelength and respective *NA*, but were not further considered in this study due to their limited availability and reproducibility (i.e. different manufacturers). Loss of quality due to a non-stationary setup (e.g. bending, vibration) was successfully solved with two feedback control loops ensuring correct coupling into the waveguide as well as maintaining best focus at long-term measurements. The use of MQTT as protocol helps to control the optical setup contact-free from multiple devices, which significantly reduces additional motion and vibrations.

With the proposed on-device image processing NN, we currently do not aim for optimized super-resolved reconstructions, but rather see it as on-device quality control (i.e. correct focusing, the variation of the light pattern) without the need of external hardware (e.g. Laptop, cloud-services). The NN also serves as a demonstration of the computational power of recent cellphones and their use in the scientific context, but remains in an experimental stage and can alternatively be replaced with the more predictable mean- or variance projection of the time-stack to reduce noise or emphasize edges.

Finally, the high refractive index waveguides used in this work are not yet available in large quantities, but will be commercially available soon. Alternative ways of creating TIRF imaging using only cover-slips [58] or optical fibers have been shown recently and are worth considering in future work. Even though we do not aim to replace commercially available SR systems with *cell*STORM II, it enables innovative ways to perform biological experiments in areas where access to these methods is limited. This includes optical detection and further study of highly pathogenic particles, such as the SARS-CoV-2 virus discovered in 2019 as we demonstrated using *d* STORM exerpiments in BSL3 laboratories. Questions such as “How does the virus react to different host cells (i.e. animal/human). How is virus entry modulated by drug treatment? Can the virus replication be influenced?”, can be potentially answered using this setup within the restricted surroundings of high-safety-level laboratories.

## 3 Methods

### 3.1 Optical Setup

The aspheric focusing lens from the OPU with a “dual-*NA*” of *NA*(*λ*_0_ = 405*nm*) = 0.9 and *NA*(*λ*_0_ = 650*nm*) = 0.5 and focal length of *f* (*λ*_0_ = 405*nm*) = 3.1*mm* can be moved in X/Z depending on the electrical current in each channel. For fluorescent excitation, we rely on different single- and multimode diode lasers (see Supp. Tab. S1.7) with excitation wavelength *λ*_*exc*_ = 445*nm*, 488*nm*, 532*nm*, 635*/*637*nm*. All parallel laser beams slightly over-fill the len’s BFP, which allows lens displacement without a serious loss of intensity in the focussed spot. By removing the focus-correcting diffractive optical element (DOE), intended to compensate the chromatic aberration, diffraction artefacts during coupling can be reduced substantially yielding increased coupling efficiency. Adding a standard cover-slip (0.15*mm*, BK7, Zeiss, Germany) between the lens and the chip reduces spherical aberration by imitating the coating material of the optical disc [32] exemplary shown in Supp. Notes S5d). Adding immersion oil as an optical matching layer applied in the gap between the WG and the coupling lens (*d*_*gap*_ ≈ 3*mm*) can help to reduce reflection losses, but leads to a destruction of the lens. The current-controlled voicecoil-mounted coupling lens is driven by a puls-width modulated signal from a microcontroller (ESP32, Espressif, China) amplified with a power transistor (BD809, Onsemiconductor, USA).

The compound microscope-based optical setup [59] (see Supp. Fig. S1.3) hosts common finite corrected objective lenses (e.g. *TL* = 160*mm*, 10×, *NA*=0.32, 40×, *NA*=0.65, 60×, *NA*=0.85 and 100×, *NA*=1.25 oil) focusable with a motorized flexure bearing-based mechanism (Fig. 1b) grey). The objective is followed by an excitation filter (see Tab. S1.7) before the intermediate image is relayed by a 10× eyepiece (Leitz, Periplan, 10 $). ce er imaging condition for common cellphone camera lenses when its exit-pupil matches the entrance pupil of the cellphone. An LED inside the chip tray for the photonic WG and inside the base-plate allows for transmission brightfield (BF) imaging by placing a standard microscope slide in the sample plane. Compared to modern infinity-corrected microscopes, the divergent beam inside the compound microscope design leads to additional aberration in fluorescent imaging since the inserted emission filter causes a defocus. By folding the the beam (new tube-length: 320*mm*) with a set of two silver-coated mirrors (Thorlabs, PF10-03-P01, 50 $) we obtain a more parallel beam path and larger magnification which results in better sampling of the PSF (i.e. smaller effective pixelsize). The PIC placed in a 3D printed tray equipped with 3 ball-magnets (neodym, 6 mm) is sitting on cylindrical screws inside a base plate (Fig. 1b) turquoise). This mechanically well-defined mechanism minimizes drift and realizes controlling its tip/tilt to adjust the optical axis with respect to the objective lens. Additionally, this mechanism allows the transport of electric power close driving LEDs for incoherent dark-field illumination imaging or heating the chip for live-cell imaging.

A pair of low-cost XY-stages (60 × 60*mm*, China, 80 $) allow a coarse alignment of the focus-spot relative to the chip (X/Y, Fig. 1b) yellow component), as well as choosing the FOV of the detection unit (Fig. 1b), red component/smartphone. Fine alignment is done by the OPU. To allow precise focusing of the objective lens, we incorporated a monolithically printed flexure bearing moved by a motorized (No Name, 28byj-48, China, 5 $) micrometer screw (0-25mm, China, 10 $) (Fig. 1b) white component). All other CAD parts are specifically designed using Inventor 2019 (Autodesk, USA) and 3D-printed (Prusa MK3S, Czech Republic) using PLA material at 100 % infill to reduce bending effects at higher temperatures (e.g. cell incubators, 37°*C*, 100% humidity, 5% *CO*_2_).

### 3.2 Software

The customized application (APP) “STORMimager” [26] running on modern Android cellphones operates the microscope’s actuators, handles scheduled frame acquisition and performs on-device image processing. This enables time-lapse imaging series (e.g. for in-vitro samples), automatic waveguide coupling, autofocusing and immediate image enhancement using neural network-based image processing of the camera stream. The compact design allows easy transportation.

All external hardware components, such as the laser, the lens or focusing motor are wired to low-cost microcontrollers (ESP32) wirelessly connected (i.e. WiFi) to the cellphone’s WiFi hotspot and communicate through the “Message Queuing Telemetry Transport” (MQTT) protocol. The protocol, where a Raspberry Pi acts as a broker (i.e. serve) and distributes messages, allows the control by multiple devices (e.g. second mobile phone). This reduces possible vibrations when manually changing parameters (i.e. coupling) during measurements is necessary since the image acquisition device is not touched. Additionally, it directly enables the creation of feedback loops from the imaging device, by changing parameters depending on a measured physical signal.

Waveguide coupling is first performed manually by coarsely moving the illumination unit with the mechanical XY-stage before the cellphone further maximizes coupling quality measured as the variance of the fluorescent or scattered signal as a function of the coupling lens position *x* (see. Supp. Notes S1.3) The software-based autofocus mechanism works similarly by moving the objective lens along the *z* (see supplement S1.2). Each optimization takes around 20 − 30 *s* and can be performed periodically during time-lapse measurements (i.e. every 30 *min*).

The code and a live-demo of the neural network is available on the website (https://beniroquai.github.io/stormocheap/). Training was performed on a GPU-equipped desktop computer (Nvidia Titan X, 6GB, Tensorflow 1.15) with an artificially created dataset based on a realistic imaging model similar to [44] (see Fig. 2 g). The newly introduced operator 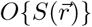 acting on the fluorophore distribution 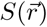 first simulates the temporally moving mode pattern *I*_*M*_ (*t*) (further described in [28]) as the sum of *N* lines at random positions 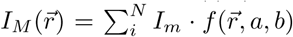 represents the intensity per mode and *f* (*x, y, b*) is a straight line *f* (*x, a, b*) = *ax* + *b* with randomly varying parameters *a* ⊂ [−*a*_*max*_*/*2, *a*_*max*_*/*2] and *b* ⊂ [2*N*_*y*_, 2*N*_*y*_]. *N*_*y*_ gives the dimension of the pixelated image along *Y* and *a*_*max*_ corresponds to a reasonable slope of the line to simulate the appearance of experimental data. The resulting illuminating field 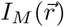 is multiplied with the downsampled (i.e. binned, *D*{·}) fluorophore distribution 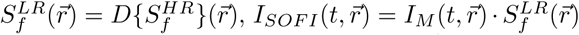, before our microscope model convolves this product with a 2D incoherent PSF (e.g. Gaussian), adds correlated Poison *P* and additive Gaussian *G* noise and simulates H.264 compression artefacts, summarized in the microscopy model operator *H*_*t*_ (further described in [16]). The result is given by (see Fig. 2 g)

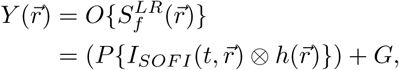

where we relied on publicly available images of microscopic samples as a source of realistic fluorophore distributions 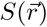 (e.g. Omero cell browser [60]) and random lines as well as dots to imitate actin networks and bacteria respectively (see example in Fig. 2c). For processing cellphone data, we used Fiji’s [49] “import image sequence” for TIF-encoded RAW images and the FFMPEG plugin (http://sites.imagej.net/Anotherche/) for MP4 video sequences. For SR reconstructions we used the SRRF [18] plugin https://github.com/HenriquesLab/NanoJ-SRRF and ThunderSTORM [52].

### 3.3 Enabling RAW Access at Video-rates

By extending the open-source camera app “FreeDCam” [51] with the module “*cell*STORM” [33], we can read-out uncompressed, unprocessed frames (no denoising, hot-pixel removal, auto white-balancing (AWB), etc.), which turn the phone used (Huawei P20 Pro) into a scientifically relevant alternative to industry-grade CMOS cameras (for further characterization see Supp. SectionS1.5). To achieve frame-rates high enough for SRI (i.e. 5 − 10 *fps*), we directly crop the incoming 12-bit byte-stream from the monochromatic camera sensor (e.g. 200 × 200 pixels) and writes it to the SD card. Even higher frame-rates (10 − 15 *fps*) can be achieved by streaming the RAW-bytes to an external server (e.g. laptop, [33]) which performs the computationally expensive process of decoding and writing the data (e.g. TIF-images).

### 3.4 Colocalization of fluorescent and antibody signal

VLP image analysis was performed using Python scripting language and custom written functions based on the program developed for [61] and employed for virus/antibody analysis in [37]. Individual viral particles were identified using an intensity maximum finding algorithm on a Gaussian smoothed image (*σ*_*pixel*_ = 2.0). Detection of maxima was kept consistent throughout using a noise tolerance parameter of 10. A circular region (⊘ = 5 pixels; 655*nm*) was then superimposed on each detected location, and all of the regions were saved for subsequent analysis. For every detected region, a random location was also generated to sample areas where eGFP/mCherry staining and thus VLPs were not likely to be present. This was achieved by randomly translating each of the detected regions to a different point within a 90-pixel radius of the original location but constrained so as not to pick an existing region, which might contain another fragment of mCherry/eGFP fluorescence. This method was effective at finding random regions that were close to virions but not overlapping and so ensured accurate comparisons between virion-containing and non-virion regions. Finally, the intensity of the antibody channel (Alexa Fluor 647) in viral and random ROIs is quantified.

### 3.5 Sample preparation

#### 3.5.1 HeLa and U2OS Cells for *d*STORM

For fixation of samples, we used the Glutaraldehyde (GA, Carl Roth, Germany) protocol from [62]. In brief, after removal of culture medium, a primary fixative of 0.4% GA and 0.25 % Triton X100 (Carl Roth, Germany) in CSB (cytoskeleton stabilizing buffer) at 37°*C* is added. After 90 *s* this solution is removed and the cells are quickly washed with 37°*C* phosphate-buffered saline (PBS, Thermo Fisher, MA, USA). The second solution of 3 % GA in Citrate-buffered saline (CBS) is then applied for 15 *min*, followed by a quick 30 *s* wash with PBS and three longer washes of 5 *min* each. To quench free aldehyde groups, a 0.3 *M* solution of glycine is then applied for 15 *min*, followed by the same washing steps as before. Actin was stained with Alexa Fluor 647-Phalloidin (Thermo Fisher), at 1:20 dilution in PBS according to the manual, by 60 min incubation at RT, before the sample was washed 3 times.

#### 3.5.2 HeLa cells for live-cell imaging

For live-cell imaging, cultured Hela cells have been transplanted to the sample container the day before imaging. SiR-Actin dye (1mM stock solution in Dimethylsulfoxid, DMSO) has been added to the cell culture medium (Dulbecco’s Modified Eagle’s Medium + 10% FBS and 1% Pen/Strep) to achieve a final concentration of 1*µM* SiR-Actin, according to [54]. Before starting the measurements, the medium has been replaced with PBS.

#### 3.5.3 Virus Isolation and Propagation (SARS-CoV-2)

SARS-CoV-2 was isolated from a respiratory sample of a patient from the Jena University Hospital (ethic approvement of the Jena University Hospital, no.: 2018-1263) and propagated by using Vero-76 cells. The virus strain was named SARS-CoV-2/hu/Germany/Jena-vi005188/2020 (5588). For virus propagation, cells were washed 12 *h* after seeding and infected with 200*νl* filtered patient sample (sterilized syringe filter, pore size 0, 2 *µl*) by the adding of Panserin 401 (PanBiotech, Germany). After five days, the cytopathic effect was detectable, and cells were frozen, centrifuged, and supernatants were obtained.

For all experiments, we used well-defined viral stocks, generated by plaque purification procedures. For this, confluent Vero-76 cell cultures were infected with serial dilutions of virus isolates diluted in EMEM for 60 *min* at 37°*C* and 5% *CO*_2_. The inoculum was exchanged with 2 *ml* MEM/BA (medium with 0.2% BSA) supplemented with 0.9% agar (Oxoid, Wesel, Germany), 0.01% DEAE-Dextran (Pharmacia Biotech, Germany) and 0.2% *NaHCO*_3_ until plaque formation was observed. Single plaques were marked using inverse microscopy. Contents of these plaques were used to infect confluent Vero-76 cell monolayers in T25 flasks. Cells were incubated at 37°*C* and 5% *CO*_2_ until pronounced cytopathic effects were visible. Then, cell cultures were frozen and clear supernatants were obtained. This plaque purification procedure was repeated. Finally, virus stocks were generated and titrated using plaque assays. For this, Vero-76 cells were seeded in 6-well plates until a 90% confluency and infected with serial dilutions of the supernatants in PBS/BA (1 *mM MgCl*_2_, 0, 9 *mM CaCl*, 0, 2%4 BSA, 100*U/ml* Pen/Strep) for 90 *min* at 37°*C*. Afterwards, cells were incubated with 2 *ml* MEM/BA (medium with 0.2% BSA) supplemented with 0.9% agar (Oxoid, Wesel, Germany), 0.01% DEAE-Dextran (Pharmacia Biotech, Germany) and 0, 2% *NaHCO*_3_ at 37° and 5% *CO*_2_ for four days. The visualization was performed by the staining with crystal violet solution (0.2% crystal violet, 20% ethanol, 3.5% formaldehyde in water) and the number of infectious particles (plaque-forming units (PFU) ml-1) was determined.

#### 3.5.4 Pseudotyped HIV Lentivirus Production

Lentiviruses were produced by transfection of the human 293T/17 cell line (ATCC catalogue# CRL-11268) with the packaging vectors pMDLg/pRRE, the PVO HIV Env expression vector and pRSV-Rev and pEGFP-Vpr. The HIV-1 Env Expression Vector (PVO, clone 4 (SVPB11) (cat# 11022, from Dr. David Montefiori and Dr. Feng Gao) [63] and pEGFP-Vpr (cat# 11386, from Dr. Warner C. Greene) [64] plasmids were obtained through the NIH AIDS Reagent Program, Division of AIDS, NIAID, NIH. Cells in a 10 *cm* dish were transfected using the calcium phosphate method with 3.5 *µg* PVO, 3.5 *µg* pMDLg/pRRE, 2.5 *µg* pRSV-Rev and 1 *µg* pEGFP-Vpr. 16h post-transfection the medium was replaced by fresh Optimem. After 48 *h* the lentiviral particles were harvested from the medium, which was passed through a 0.45 *µm* PES filter and concentrated by ultracentrifugation at 76, 000 × *g* (avg.) through a 20% sucrose cushion. The pellet was gently resuspended in PBS. Viruses were snap-frozen and kept at −80° *C*.

#### 3.5.5 Production of Lentivirus VLPs pseudotyped with SARS-CoV-2 Spike Proteins

Lentivirus-derived VLPs were produced by transfecting 10 cm dishes of HEK293T LentiX cells with lentiviral packaging plasmids encoding Gag/Pol (pMDLg/pRRE, 15 *µg*) and Rev (pRSV-REV, 6 *µg*) (Addgene plasmids 12251 and 12253) together with vectors expressing SARS-CoV-2 Spike (13 *µg*) and MyrPalm-mCherry (5 *µg*) using Lipofectamin 2000. The plasmids encoding the Spike protein were a mixture of (10 *µg*) of pcD*NA*3.1-SARS2-Spike (Addgene plasmid 145032) and (3 *µg*) of GFP-tagged S protein (Genscript, MC 0101089). 48 *h* after transfection, the supernatant was collected, cleared by filtering through a 0.45 *µm* filter and concentrated by ultracentrifugation through a 20% sucrose cushion. VLPs were resuspended in Life cell imaging solution (Thermo Fisher) and stored at − 80°*C*. Formation of VLPs was verified by electron microscopy.

#### 3.5.6 Immunostaining of Active and pseudotpyed SARS-CoV-2

Active and pseudotyped SARS-CoV-2 particles were immobilized by adding 0.01% Poly-L-Lysine (PLL, Sigma Aldrich, MIS, USA) for 30 *min*, before the surface was rinsed with *H*_2_*O*. Concentrated virus stock (5 − 10 *µL*) were placed on the surface and incubated for 30 *min* at room temperature (RT). Blocking was performed with 4% BSA for 60 *min*. For active and pseudotyped SARS-CoV-2 particles, we add primary 1:200 *α*-spike antibody (1A9, Genetex, CL, USA) in BSA overnight at 7°*C*. Sample was washed three times with PBS, before secondary 1:200 *α*-mouse antibody pre-labelled with Alexa Fluor 647 (*AB*_2_535804, Thermofisher, MA, USA) in BSA were incubated at RT. Sample was washed 3× with BSA before imaging. For *d* STORM measurements OxEA was used as the imaging buffer. In case of HIV VLPs 10E8 (produced in bacteria, a kind gift of Prof. José L Nieva) was incubated at 1 *µM* concentration in 2% BSA/PBS for 1 *h*, and revealed with a secondary anti-human IgG labelled with Alexa Fluor 647 (1:250) for 1 *h*.

#### 3.5.7 *E. coli* Bacteria

To a suspension of living *E. coli* (DH5-alpha-lambda-pir.-coli) in LB-medium we added 0.4*M* mCLING Atto647N (Synaptic system, Goettingen, Germany) and incubated it for 5 minutes. The concentration of the bacteria was achieved using ultra-centrifuging at 10:000 rounds/min for 3 minutes. After aspirating the top layer, 4% PFA is added. Removing unbound dyes is done by repeatedly centrifuging, aspirating the top layer and refilling it with PBS. Coverslips are prepared by adding 0.01% PLL (Sigma Aldrich) to the surface and incubate it for 30 *min* RT. After removing the PLL, the bacteria were added and incubated for another 30 *min*. We washed it 3× using PBS and sealed it on a microscope cover glass. This preparation was also used on the surface of waveguide chips, where the sample buffer (PBS) is contained inside a Polydimethylsiloxane (PDMS) sample chamber, and sealed with a coverslip.

## Supporting information

Supp. Video 2: Screencast of NN-based SOFI

Supp. Video 3: dSTORM using cellSTORM II

Supp. Video 3: SOFI using cellSTORM II

Supp. Video 4: Time-lapse of in-vitro HeLa cells

## Authors Contribution

**Conceptualization:** B.D., Ø.H.

**Data curation:** B.D.

**Formal analysis:** B.D., Ø.H., P.T., K.O.S., S.D.-E., P.C., R.H., B.S.

**Funding acquisition:** B.D., B.S., R.H., C.E.

**Investigation:** B.D.,Ø.H., F.H., K.O.S., S.C.

**Methodology:** B.D., B.S.

**Project administration:** B.D.

**Resources:** B.D., R.L., S.C., A.M., C.E.

**Software:** B.D.

**Supervision:** B.D.

**Validation:** B.D., Ø.H., P.T., K.O.S., S.D.-E., P.C., R.H., B.S.

**Visualization:** B.D.

**Writing – original draft:** B.D., Ø.H., P.T., C.E., with input from all authors.

**Writing – review & editing:** All authors

## Acknowledgements

B.S.A. acknowledges funding from European Commission ERC Starting Grant (336716) and ERC PoC 957464 and), Research Council of Norway (Grant 288565). We acknowledge funding by the DFG Transregio Project TRR166, TP04 (PT) and the Leibniz ScienceCampus InfectoOptics SAS-2015-HKI-LWC (AJ). The authors further acknowledge the support of this work by a grant from the IZKF (ACSP02) (SDE). The study was further funded by the Deutsche Forschungsgemeinschaft (DFG, German Research Foundation) under Germany’s Excellence Strategy (EXC 2051, Project-ID 390713860). This work was financially supported by the Deutsche Forschungsgemeinschaft through the Cluster of Excellence “Balance of the Microverse” under Germany’s Excellence Strategy – EXC 2051 – Project-ID 690 390713860. We thank Fatina Siwczak and Swen Carlstedt for fruitful discussions. We further thank Merete Storflor and Sören Abel (Infection Biology Research Group, UiT) for providing the *E.coli* samples and Deanna Wolfson (Optics group, UiT) for help with preparation of the *E.coli* samples. We would like to thank Sindy Burgold-Voigt and Ralf Ehricht from Leibniz IPHT Jena for their great methodical contribution and the fruitful discussions. We thank especially Ingo Fuchs who helped a lot in understanding the principles of the acquisition process of cellphone cameras as well as his support of the software design. Pablo Carravilla acknowledges the Basque Government postdoctoral program (POS_2019_2_0022) for funding his position. pMDLg/pRRE and pRSV-Rev were a gift from Didier Trono (Addgene plasmid #12251 and #12253; http://n2t.net/addgene:12251 and http://n2t.net/addgene:12253; RRID:Addgene_12251 and RRID:Addgene_12253). K.O.S is supported by a Career fellowship from the South-Eastern Norway Regional Health Authority.

## Conflicting Statements

B.S.A. has applied for two patents for chip-based optical nanoscopy. B.S.A. and O.I.H. are co-founder of the company Chip NanoImaging AS, which commercializes on-chip super-resolution microscopy systems.

## References

[1] Marco Fritzsche and Guillaume Charras. “Dissecting protein reaction dynamics in living cells by fluorescence recovery after photobleaching”. In: Nature Protocols (2015). ISSN: 17502799.

[2] Knut Rennert et al. “A microfluidically perfused three dimensional human liver model”. In: Biomaterials 71 (Dec. 2015), pp. 119–131. ISSN: 0142-9612. URL: https://www.sciencedirect.com/science/article/pii/S0142961215007097.

[3] Marko Gröger et al. “Monocyte-induced recovery of inflammation-associated hepatocellular dysfunction in a biochip-based human liver model”. In: Scientific Reports 6.1 (Apr. 2016), p. 21868. ISSN: 2045-2322. URL: http://www.nature.com/articles/srep21868.

[4] Brandon Berg et al. “Cellphone-Based Hand-Held Microplate Reader for Point-of-Care Testing of Enzyme-Linked Immunosorbent Assays”. In: ACS Nano (2015). ISSN: 1936086X.

[5] Jakub Chojnacki and Christian Eggeling. Super-resolution fluorescence microscopy studies of human immunodeficiency virus. 2018.

[6] Ernst Abbe. “Beiträge zur Theorie des Mikroskops und der mikroskopischen Wahrnehmung”. In: Archiv für mikroskopische Anatomie 9.1 (1873), pp. 413–418. ISSN: 0176-7364. URL: http://dx.doi.org/10.1007/BF02956173.

[7] S W Hell and J Wichmann. “Stimulated emission depletion fluorescence microscopy”. In: Optics letters (1994). ISSN: 0146-9592.

[8] Rainer Heintzmann and Mats G. L. Gustafsson. “Subdiffraction resolution in continuous samples”. In: Nature Photonics 3.7 (2009), pp. 362–364. ISSN: 1749-4885. URL: http://dx.doi.org/10.1038/nphoton.2009.102.

[9] Eric Betzig et al. “Imaging intracellular fluorescent proteins at nanometer resolution.” In: Science (New York, N.Y.) 313.5793 (2006), pp. 1642–5. ISSN: 1095-9203. URL: http://www.ncbi.nlm.nih.gov/pubmed/16902090.

[10] Jianquan Xu, Hongqiang Ma, and Yang Liu. “Stochastic optical reconstruction microscopy (STORM)”. In: Current Protocols in Cytometry (2017). ISSN: 19349300.

[11] Sebastian van de Linde et al. “Direct stochastic optical reconstruction microscopy with standard fluorescent probes.” In: Nature protocols 6.7 (2011), pp. 991–1009. ISSN: 1750-2799. URL: http://www.ncbi.nlm.nih.gov/pubmed/21720313.

[12] Hongqiang Ma et al. “A simple and cost-effective setup for super-resolution localization microscopy”. In: 7.1 (2017), p. 1542. ISSN: 2045-2322. URL: http://www.ncbi.nlm.nih.gov/pubmed/28484239%20http://www.pubmedcentral.nih.gov/articlerender.fcgi?artid=PMC5431525%20http://www.nature.com/articles/s41598-017-01606-6.

[13] Kwasi Kwakwa et al. “easySTORM: a robust, lower-cost approach to localisation and TIRF microscopy”. In: Journal of Biophotonics 9.9 (2016), pp. 948–957. ISSN: 18640648.

[14] Thorge Holm et al. “A Blueprint for Cost-Efficient Localization Microscopy”. In: ChemPhysChem 15.4 (2014), pp. 651–654. ISSN: 14394235. URL: http://doi.wiley.com/10.1002/cphc.201300739%20http://onlinelibrary.wiley.com/doi/10.1002/cphc.201300739/abstract.

[15] Robin Diekmann et al. “Characterization of an industry-grade CMOS camera well suited for single molecule localization microscopy – high performance super-resolution at low cost”. In: Scientific Reports 7.1 (Dec. 2017), p. 14425. ISSN: 2045-2322. URL: http://www.nature.com/articles/s41598-017-14762-6www.nature.com/scientificreports.

[16] Benedict Diederich et al. “cellSTORM—Cost-effective super-resolution on a cellphone using dSTORM”. In: PLOS ONE 14.1 (Jan. 2019). Ed. by Tom Waigh, e0209827. ISSN: 1932-6203. URL: http://dx.plos.org/10.1371/journal.pone.0209827.

[17] Martin Weigert et al. “Content-Aware Image Restoration: Pushing the Limits of Fluorescence Microscopy”. In: (2017). URL: http://dx.doi.org/10.1101/236463.

[18] Nils Gustafsson et al. “Fast live-cell conventional fluorophore nanoscopy with ImageJ through super-resolution radial fluctuations”. In: Nature Communications (2016). ISSN: 20411723.

[19] Elias Nehme et al. “Deep-STORM: Super Resolution Single Molecule Microscopy by Deep Learning”. In: (Jan. 2018). 1801.09631. URL: http://arxiv.org/abs/1801.09631.

[20] Nicholas Boyd et al. “DeepLoco: Fast 3D Localization Microscopy Using Neural Networks”. In: bioRxiv (Feb. 2018), p. 267096. URL: https://www.biorxiv.org/content/early/2018/02/16/267096.

[21] Idir Yahiatene et al. “Entropy-Based Super-Resolution Imaging (ESI): From Disorder to Fine Detail”. In: ACS Photonics (2015). ISSN: 23304022.

[22] Wei Ouyang et al. “Deep learning massively accelerates super-resolution localization microscopy”. In: Nature Biotechnology (2018). ISSN: 15461696.

[23] James P. Sharkey et al. “A one-piece 3D printed flexure translation stage for open-source microscopy”. In: Review of Scientific Instruments 87.2 (Feb. 2016), pp. 1–7. ISSN: 10897623. 1509.05394. URL: http://scitation.aip.org/content/aip/journal/rsi/87/2/10.1063/1.4941068.

[24] James W.P. Brown et al. “Single-molecule detection on a portable 3D-printed microscope”. In: Nature Communications (2019). ISSN: 20411723.

[25] Koen J.A. Martens et al. “Visualisation of dCas9 target search in vivo using an openmicroscopy framework”. In: Nature Communications (2019). ISSN: 20411723.

[26] Benedict Diederich et al. “UC2 – A Versatile and Customizable low-cost 3D-printed Optical Open-Standard for microscopic imaging”. In: bioRxiv (Jan. 2020), p. 2020.03.02.973073. URL: http://biorxiv.org/content/early/2020/03/03/2020.03.02.973073.abstract.

[27] Robin Diekmann et al. “Chip-based wide field-of-view nanoscopy”. In: Nature Photonics 11.5 (2017), pp. 322–328. ISSN: 17494893.

[28] Øystein I Helle et al. Structured illumination microscopy using a photonic chip. Tech. rep. 1903.05512v1. URL: https://arxiv.org/pdf/1903.05512.pdf.

[29] Fabrizio La Rosa et al. “Optical Image Stabilization”. In: Figure 1 (2011), pp. 1–26. URL: http://www.st.com/web/en/resource/technical/document/white%7B%5C_%7Dpaper/ois%7B%5C_%7Dwhite%7B%5C_%7Dpaper.pdf.

[30] Tigran Galstian. Smart mini-cameras. 2006, p. 323. ISBN: 9781466512924.

[31] L. K. Lai and T. S. Liu. “Design of auto-focusing modules in cell phone cameras”. In: International Journal on Smart Sensing and Intelligent Systems 4.4 (2011), pp. 568–582. ISSN: 11785608.

[32] Edwin En Te Hwu and Anja Boisen. Hacking CD/DVD/Blu-ray for Biosensing. 2018.

[33] Benedict Diederich. Project Page STORM-on-the-chea(i)p. 2020. URL: https://beniroquai.github.io/stormocheap/ (visited on 05/16/2020).

[34] Nikhil Jayakumar et al. “On-chip TIRF nanoscopy by applying Haar wavelet kernel analysis on intensity fluctuations induced by chip illumination”. In: (July 2020). 2007.12899. URL: http://arxiv.org/abs/2007.12899.

[35] T Dertinger et al. “fluctuation imaging (SOFI)”. In: Proceedings of the National Academy of Sciences of the United States of America (2009). ISSN: 1091-6490.

[36] T. Dertinger et al. “Fast, background-free, 3D super-resolution optical fluctuation imaging (SOFI)”. In: Proceedings of the National Academy of Sciences of the United States of America (2009). ISSN: 00278424.

[37] Andrey Ignatov et al. “AI Benchmark: Running deep neural networks on android smart-phones”. In: Lecture Notes in Computer Science (including subseries Lecture Notes in Artificial Intelligence and Lecture Notes in Bioinformatics). 2019. ISBN: 9783030110208. 1810.01109.

[38] Martín Abadi et al. “TensorFlow: Large-Scale Machine Learning on Heterogeneous Distributed Systems”. In: (Mar. 2016). 1603.04467. URL: http://arxiv.org/abs/1603.04467.

[39] Sepp Hochreiter and Jürgen Schmidhuber. “Long Short-Term Memory”. In: Neural Computation (1997). ISSN: 08997667.

[40] Xingjian Shi et al. “Convolutional LSTM network: A machine learning approach for precipitation nowcasting”. In: Advances in Neural Information Processing Systems. 2015. 1506.04214.

[41] Seyed Shahabeddin Nabavi, Mrigank Rochan, and Yang Wang. Future Semantic Segmentation with Convolutional LSTM. Tech. rep. 1807.07946v1. URL: https://arxiv.org/pdf/1807.07946v1.pdf.

[42] B. E. a. Saleh and M. C. Teich. Grundlagen der Photonik. 2008, p. 1406. ISBN: 9783527406777. URL: http://books.google.com/books?hl=en%7B%5C&%7Dlr=%7B%5C&%7Did=dXSfqi1izUkC%7B%5C&%7Doi=fnd%7B%5C&%7Dpg=PA1%7B%5C&%7Ddq=Grundlagen+der+Photonik%7B%5C%7Dots=J%7B%5C_%7D5%7B%5C_%7D5wrjEB%7B%5C%7Dsig=wKTKYvIgCo5s74Htvv3LoDA%7B%5C_%7D1Sw.

[43] Wenzhe Shi et al. “Real-Time Single Image and Video Super-Resolution Using an Efficient Sub-Pixel Convolutional Neural Network”. In: Proceedings of the IEEE Computer Society Conference on Computer Vision and Pattern Recognition. 2016. ISBN: 9781467388504. 1609.05158.

[44] Charles N. Christensen et al. “ML-SIM: A deep neural network for reconstruction of structured illumination microscopy images”. In: (Mar. 2020). 2003.11064. URL: http://arxiv.org/abs/2003.11064.

[45] Carolin Vietz et al. “Benchmarking Smartphone Fluorescence-Based Microscopy with DNA Origami Nanobeads: Reducing the Gap toward Single-Molecule Sensitivity”. In: ACS Omega 4.1 (Jan. 2019), pp. 637–642. ISSN: 2470-1343. URL: http://pubs.acs.org/doi/10.1021/acsomega.8b03136.

[46] ITU-T. “H.264”. In: International Telecommunication Union (2013).

[47] Jürgen Popp et al., eds. Handbook of Biophotonics. Weinheim, Germany: Wiley-VCH Verlag GmbH & Co. KGaA, Jan. 2013. URL: http://doi.wiley.com/10.1002/9783527643981.

[48] Niccolò Banterle et al. “Fourier ring correlation as a resolution criterion for super-resolution microscopy”. In: Journal of Structural Biology 183.3 (2013), pp. 363–367. ISSN: 10478477.

[49] Johannes Schindelin et al. Fiji: An open-source platform for biological-image analysis. 2012.

[50] Leila Nahidiazar et al. “Optimizing imaging conditions for demanding multi-color super resolution localization microscopy”. In: PLoS ONE (2016). ISSN: 19326203.

[51] Benedict Diederich and Ingo Fuchs. Github: cellSTORM Android APP. 2018. URL: https://github.com/bionanoimaging/cellSTORM-ANDROID.

[52] Martin Ovesný et al. “ThunderSTORM: a comprehensive ImageJ plug-in for PALM and STORM data analysis and super-resolution imaging”. In: Bioinformatics 30.16 (Aug. 2014), pp. 2389–2390. ISSN: 1367-4803. URL: http://www.ncbi.nlm.nih.gov/pubmed/24771516%20http://www.pubmedcentral.nih.gov/articlerender.fcgi?artid=PMC4207427%20https://academic.oup.com/bioinformatics/article-lookup/doi/10.1093/bioinformatics/btu202.

[53] Romain F. Laine et al. “NanoJ: A high-performance open-source super-resolution microscopy toolbox”. In: Journal of Physics D: Applied Physics (2019). ISSN: 13616463.

[54] Gražvydas Lukinavičius et al. “Fluorogenic probes for live-cell imaging of the cytoskeleton”. In: Nature Methods 11.7 (2014), pp. 731–733. ISSN: 15487105.

[55] Xiong Ding et al. Interfacing Pathogen Detection with Smartphones for Point-of-Care Applications. 2019.

[56] Pablo Carravilla et al. “Molecular recognition of the native HIV-1 MPER revealed by STED microscopy of single virions”. In: Nature Communications (2019). ISSN: 20411723.

[57] Ronald L Gordon. “Exact computation of scalar 2D aerial imagery”. In: Design, Process Integration, and Characterization for Microelectronics 4692.1 (2002), pp. 517–528. URL: http://link.aip.org/link/?PSI/4692/517/1.

[58] Yulung Sung, Fernando Campa, and Wei-Chuan Shih. “Open-source do-it-yourself multicolor fluorescence smartphone microscopy”. In: Biomedical Optics Express 8.11 (Nov. 2017), p. 5075. ISSN: 2156-7085. URL: https://www.osapublishing.org/abstract.cfm?URI=boe-8-11-5075.

[59] Zachary F. Phillips et al. “Multi-Contrast Imaging and Digital Refocusing on a Mobile Microscope with a Domed LED Array”. In: Plos One 10.5 (May 2015). Ed. by Jonathan A Coles, e0124938. ISSN: 1932-6203. URL: http://dx.plos.org/10.1371/journal.pone.0124938%20http://www.ncbi.nlm.nih.gov/pubmed/25969980%20http://www.pubmedcentral.nih.gov/articlerender.fcgi?artid=PMC4430423.

[60] Chris Allan et al. OMERO: Flexible, model-driven data management for experimental biology. 2012.

[61] Silvia Galiani et al. “Super-resolution microscopy reveals compartmentalization of peroxisomal membrane proteins”. In: Journal of Biological Chemistry (2016). ISSN: 1083351X.

[62] Donna R. Whelan and Toby D.M. Bell. “Image artifacts in single molecule localization microscopy: Why optimization of sample preparation protocols matters”. In: Scientific Reports 5 (2015). ISSN: 20452322.

[63] Ming Li et al. “Human Immunodeficiency Virus Type 1 env Clones from Acute and Early Subtype B Infections for Standardized Assessments of Vaccine-Elicited Neutralizing Antibodies”. In: Journal of Virology (2005). ISSN: 0022-538X.

[64] Evelyne Schaeffer, Romas Geleziunas, and Warner C. Greene. “Human Immunodeficiency Virus Type 1 Nef Functions at the Level of Virus Entry by Enhancing Cytoplasmic Delivery of Virions”. In: Journal of Virology (2001). ISSN: 0022-538X.

