## Supplementary Information for "Nanoscopy on the Chea(i)p"

### Contents


S1 Supplementary Image . . . . . . . . . . S3
- S1.1 Reconstruction artifacts using compressed video data in SRRF . . . . . . S3
- S1.2 Comparison dSTORM RAW vs MP4 . . . . . . S4
- S1.3 Optical Setup of *cell*STORM . . . . . . S5

S1 Supplementary Video . . . . . . . . . . S6
- S1.1 Long-term time-series of Actin-labelled HeLa cells . . . . . . S6
- S1.2 Screencast of On-device Super-resolution Imaging . . . . . . S6
- S1.3 *d*STORM experiment using cellSTORM II . . . . . . S6
- S1.4 SOFI experiment using cellSTORM II . . . . . . S6

S1 Supplementary Notes . . . . . . . . . . S7
- S1.1 Bill of Materials . . . . . . S7
- S1.2 Algorithm for the Autofocus . . . . . . S9
- S1.3 Algorithm for the automatic coupling . . . . . . S11
- S1.4 Build instructions . . . . . . S13
- S1.5 ISO vs Noise on P20 . . . . . . S13
- S1.6 Converting any microscope into a WG-based microscope . . . . . . S15
- S1.7 Dual-colour mode for co-localization . . . . . . S16
- S1.8 Effective Pixelsizes . . . . . . S18

### S1 Supplementary Image

#### S1.1 Reconstruction artifacts using compressed video data in SRRF

The fourier-based image reconstruction method SRRF shows imaging artifacts on the compression grid of the H264-based algorithm.

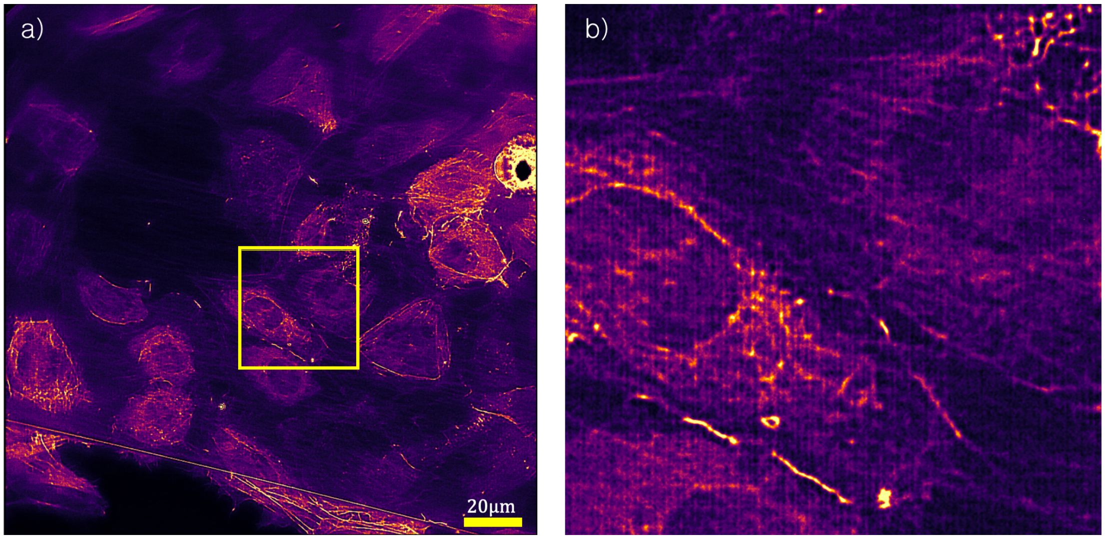


**Figure S1. Reconstruction artifacts using SRRF in MP4** - The integer tansform based compression algorithm H.264 promotes discontinuities at the border of unit cell. Common reconstruction algorithms like SRRF [1] lead to artifacts at these edges.

#### S1.2 Comparison dSTORM RAW vs MP4



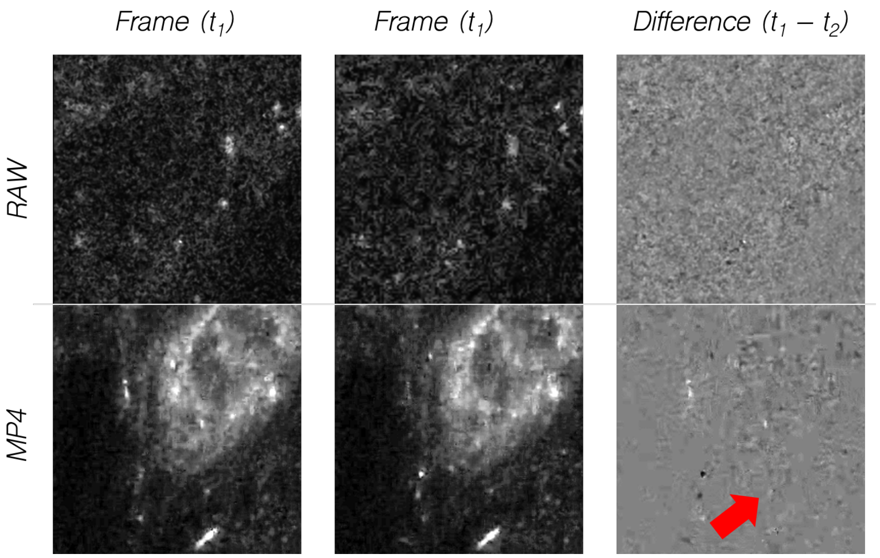


**Figure S2. Comparison between *d*STORM experiments using RAW and MP4 -** We acquire a similar sample (HeLa) labelled with Alexa Fluor 647 under the same experimental condition (i.e. imagin buffer/MEA, intensity). By computing the difference between two adjacent frames from a *d*STORM experiment, It is directly apparent, that the compression algorithm (MP4) destroys the noise characteristic. Noise is therefore no longer correlated. Additionally, the MP4 compressed data shows cutting edges the noise. The RAW-frame acquisition shows no cutting edges.

#### S1.3 Optical Setup of *cell*STORM

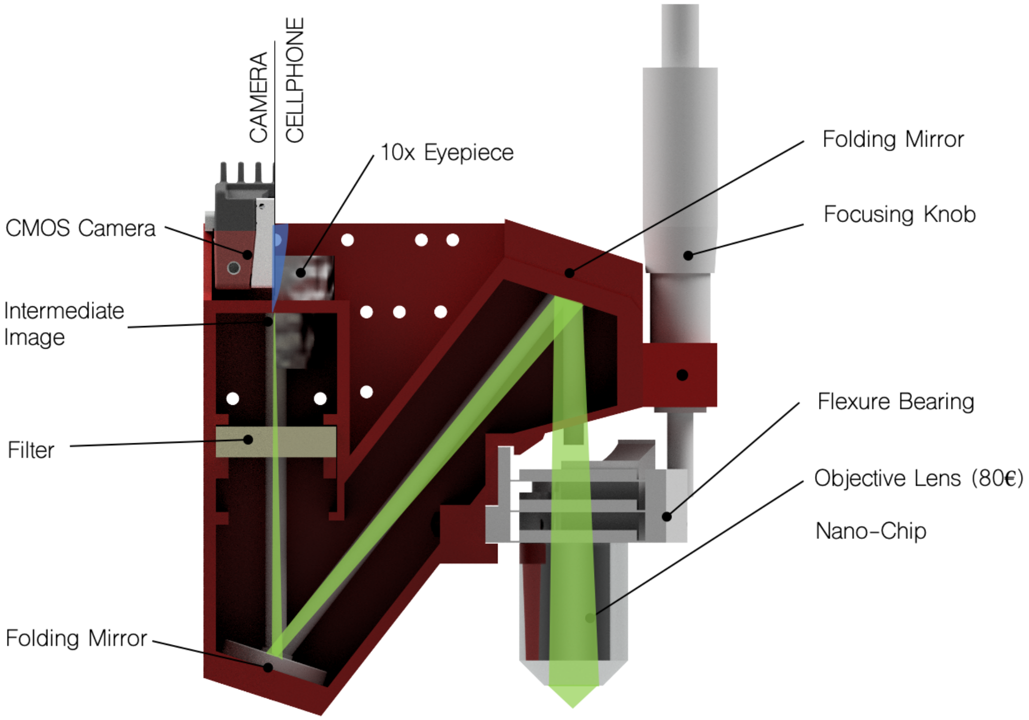


**Figure S3. Optical Setup of *cell*STORM II -** The optical setup is deviated from a classical finite-corrected microscope design which hosts a finite-corrected objective lens with an intermediate image to be found at a distance of the tube length (green beam-path). In order to reduce spherical aberration due to the non-parallel beam path when inserting the emission filter, the tube-length is elongated from 160 $mm$ to 320 $mm$. The intermediate image is relayed by an eyepiece in order to image it with the cellphone camera or directly imaged with an industry-grade CMOS camera (e.g. Allied Vision Alvium 1800 U-158)

### S1 Supplementary Video

#### S1.1 Long-term time-series of Actin-labelled HeLa cells

The video shows a $17h$ time-lapse of SIR-rhodamine labelled HeLa cells at room-temperature. Images were taken every $4\,min$.

#### S1.2 Screencast of On-device Super-resolution Imaging

Using the APP "STORMImager", the processed screen overlay of the cropped centre region of the video stream shows the result of the NN-based image processing algorithm. In case of the experiment while the excitation intensity is randomly fluctuating, the membrane of *E. Coli* bacteria can be recovered at a frame rate of $1-2\,fps$.

#### S1.3 *d*STORM experiment using cellSTORM II

A common *d*STORM experiment of Alexa Fluor Phal. labelled HeLa cells. The setup experiences a temporal drift which can be compensated in post-processing. The video was acquired with the Huawei P20 Pro using the monochrome camera. The APP FreedCam recorded the video-stream at $t_{exp} = 1/20\,s$ and $ISO = 3200$.

#### S1.4 SOFI experiment using cellSTORM II

Fluctuating the coupling lens along Z causes a temporally varying mode pattern inside the waveguide. The varying excitation pattern and resulting changing change in fluorescence response can be used for super-resolution imaging suitable for living samples since the excitation intensity can be relatively low. The video shows fixed *E. Coli* labelled with mCLING-ATTO647.

### S1 Supplementary Notes

#### S1.1 Bill of Materials

| Part | Name | Price | Supplier |
|---|---|---|---|
| Laser (red) | Laserlands # 3450, $P = 300\,mW$ $\lambda_0 = 637\,nm$, Dot Laser Module TTL/analog $12\,V$ DC | 50 € | Laserlands |
| Laser (green) | $P = 200\,mW$ $\lambda_0 = 532\,nm$, Dot Laser Module TTL/analog $12\,V$ DC | 90 € | lilly electronics |
| Objective Lens | BRESSER DIN-Objektiv 60x, NA 0.85, 160/0.17 | 45 € | Ebay |
| 2x Mirror | PF10-03-P01 diameter:1" Protected Silver Mirror | 50 € | Thorlabs |
| 2x XY-Stages | XY Axis Manual Trimming Platform Linear Stage Tuning Sliding Table $60 \times 60\,mm$ | 80 € | Ebay |
| Longpass Filter (red) | Chroma, HQ675/50m, 25mm diameter | 200 € | Chroma |
| Eyepiece | 10× Periplan Leitz Wetzlar, Germany | 18 € | Ebay |
| ESP32 | 3x ESP32 Dev Kit, WROOM32, NodeMCU | 7 € | Ebay |
| LED Buk driver | 3x SPARKFUN ELECTRONICS INC. COM-13705 | 18 € | Sparkfun |
| Wires | Various | 10 € | Ebay |
| Screws | M3/M4 cylinder head, galvanized, 16/25/30$mm$ | 10 € | Würth |

| Part | Name | Price | Supplier | |
|---|---|---|---|---|
| Powersupply | 5V, 3A, USB | 10 € | Ebay | |
| MQTT Broker | Raspberry Pi v3B, SD-Card (16 GB) + 5V Powersupply | 70 € | Ebay | |
| Optical Pickup | Sony KES-400A (Playstation 3, Sony) | 1-10 € | Ebay | |
| PLA filament | Prusament, 1.75 $mm$ black, 1 $Kg$ | 30 € | Prusa | 48 |
| Cellphone | Huawei P20 Pro | 300 € | Ebay | |
| Ball Magnets | Neodym Magnets, 5 $mm$ diameter | 4 € | Ebay | |
| Micrometer Screw | 30 $mm$ micrometer screw | 15 € | Ebay | |
| Stepper Motor | 28byj-48, Driver Board | 5 € | Ebay | |

#### S1.2 Algorithm for the Autofocus

We implemented a basic autofocus algorithm to compensate mechanical drift in long-term experiments. The mechanism (Fig. S4 a/b) is based on a micro-meter screw (MS) which pushes a flexure bearing (FB). The mechanically reduced ($m \approx 2$) level arm moves the objective arm up and down in order to refocus the sample. A stepper motor (M) (28byj-48, no name, China), is connected to the MS using a gear belt. The autofocusing mechanism as indicated in Fig. S4

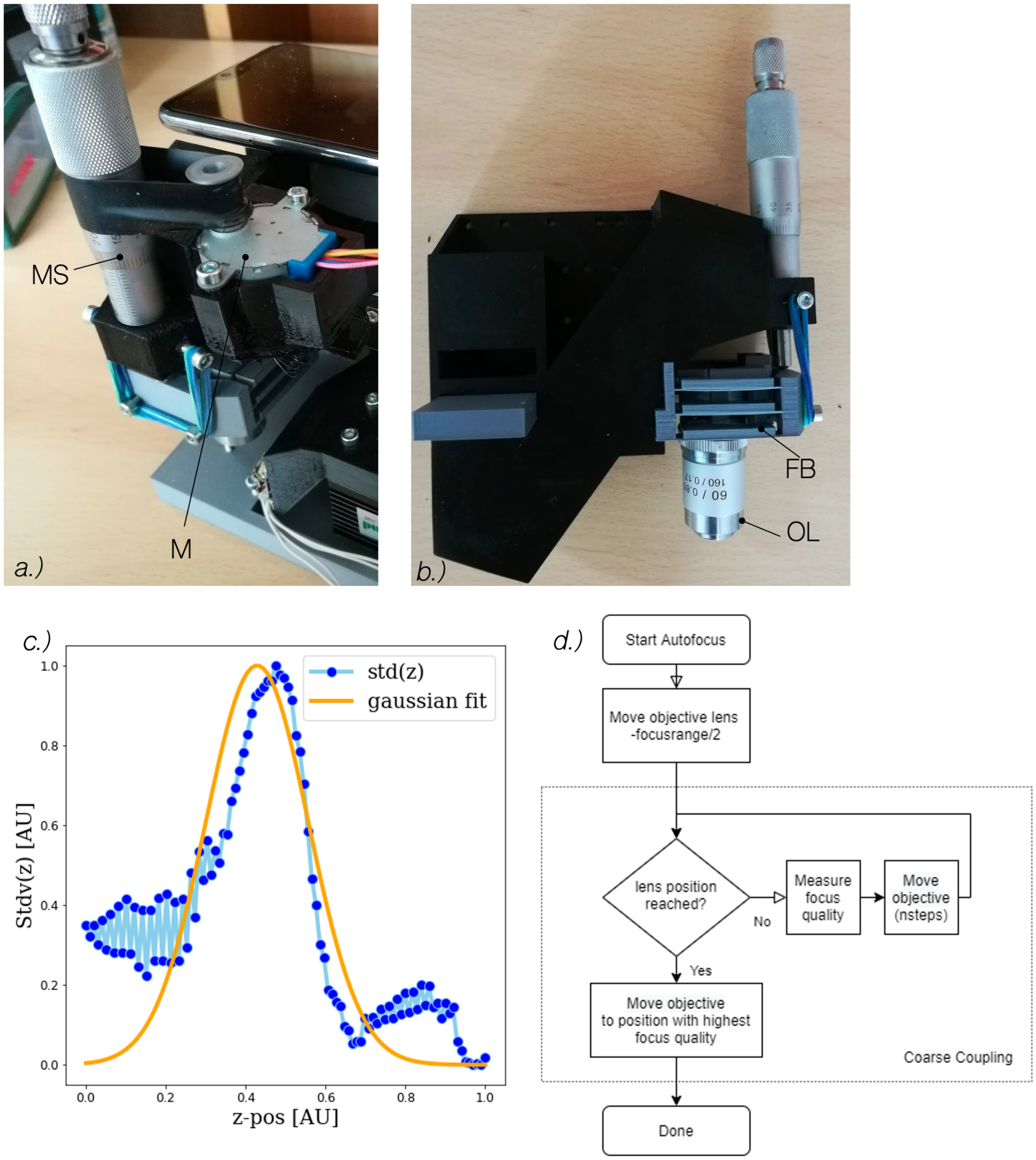


**Figure S4. Flowchart of the autofocus algorithm** - A basic hardware add-on mounts a stepper-motor (M) to the assembly a) which turns the micrometer screw (MS). The objective lens (OL) mounted on the flexure bearing (FB) moves up and down by pushing it using the MS. A basic autofocus algorithm d) finds the best z-position which maximizes the image contrast, exemplary plotted in c)

looks for the highest contrast as a function of the z-coordinate

$$\underset{\rightarrow z}{argmax}\; f_{focus}(z) \tag{1}$$

$$\text{where,} \tag{2}$$

$$f(z) = var(I(z)). \tag{3}$$

The software-based autofocus algorithm assumes, that an intensity signal from a scattering or fluorescent source is detected by the camera. A focus scan usually takes between $30 - 60\,s$ depending on the present focus range. Focussing and image acquisition at the same time is possible, but not yet implemented.

#### S1.3 Algorithm for the automatic coupling

Similar to the autofocus mechanism in Sec. S1.2, we implement an automatic coupling procedure. The aim of this algorithm is to maximize and maintain coupling efficiency by creating a feedback loop-based on the observed variance in an image (i.e. fluorescence, scattered signal) as a function of the coupling lens position.
The OPU lens visualized in Fig. S5 is equipped with two voice coil motors (VCM) for moving the aspheric lens ($NA(\lambda = 405\,nm) = 0.85,\ f' = 3.1\,mm$) in x- and z-direction. We choose to take the OPU from an used Playstation 3 (KES400-A, [2, 3, 4]) drive since it is inexpensive and widely available to maintain reproducibility. The lens, can accurately be controlled using a PWM signal from an amplified output of a microcontroller. It is of great importance to have a high carrier frequency in the PWM signal to avoid unstable coupling. At the same time, a high resolution is required in order to move the lens with sub-micrometer precision. We choose a resolution of 15Bit, which divides the scan-range of about $1.5\,mm$ into $\approx 45\,nm$/step. A detailed description to replicate this setup using off-the-shelf components is provided in our project-webpage. Different OPU lenses with lower numerical apertures can lead to an increased coupling efficiency.
During the experiments, we saw an increase in coupling efficiency, when the diffractive optical element mounted in the back of the lens was removed. It causes unwanted interference effects at the air-chip boundary. Using immersion oil at the air-lens interface as an optical bonding material can lead to a destruction of the hollow aspheric lens. Adding a cover slip (BK7, $0.135\,\mu m$, Zeiss, Germany) between the lens (see Fig. S5 c) compensates the pre-compensated spherical aberration of the OPU lens, further quantified with an optical simulation in ZEMAX (MA, USA). A similar model of a DVD OPU was used to demonstrate the decreased coupling efficiency while varying the thickness of the cover material (Fig. S5 d/e).

The automatic coupling mechanism visualized in Fig. S5 b) is divided into two stages, where first a coarse alignment is performed, which is followed by a fine adjustment. The algorithm requires a previous manual coupling with already good coupling efficiency using the XY-stage. In order to simplify the algorithm we neglect lens movement in z-direction. Similar to the autofocus system mentioned above, the cellphone observes the fluorescence/scattered signal perpendicular to the waveguide and measures the coupling quality $f_{coupling}$ as a function of the lens position in $x$-direction $p_x$. Initially, the cellphone sends control commands to the MQTT-connected OPU with a coarse stepsize (e.g. $4.5\,\mu m$/step) and sweeps through the full scan range. The maximum of this procedure is the centre position of the second "fine coupling" with smallest stepsize (e.g. $45\,nm$/step). The goal for the automatic coupling routine is to find the highest efficiency given as

$$\underset{\rightarrow p_x}{argmax} f_{coupling}(z) \tag{4}$$

$$\text{where,} \tag{5}$$

$$f_{coupling}(z) = var(I(x_p)). \tag{6}$$

The calculation is carried out using the open source library OpenCV [5] inside the Android APP "STORMImager" available on our project-webpage and runs at $\approx 1\,s$/step.

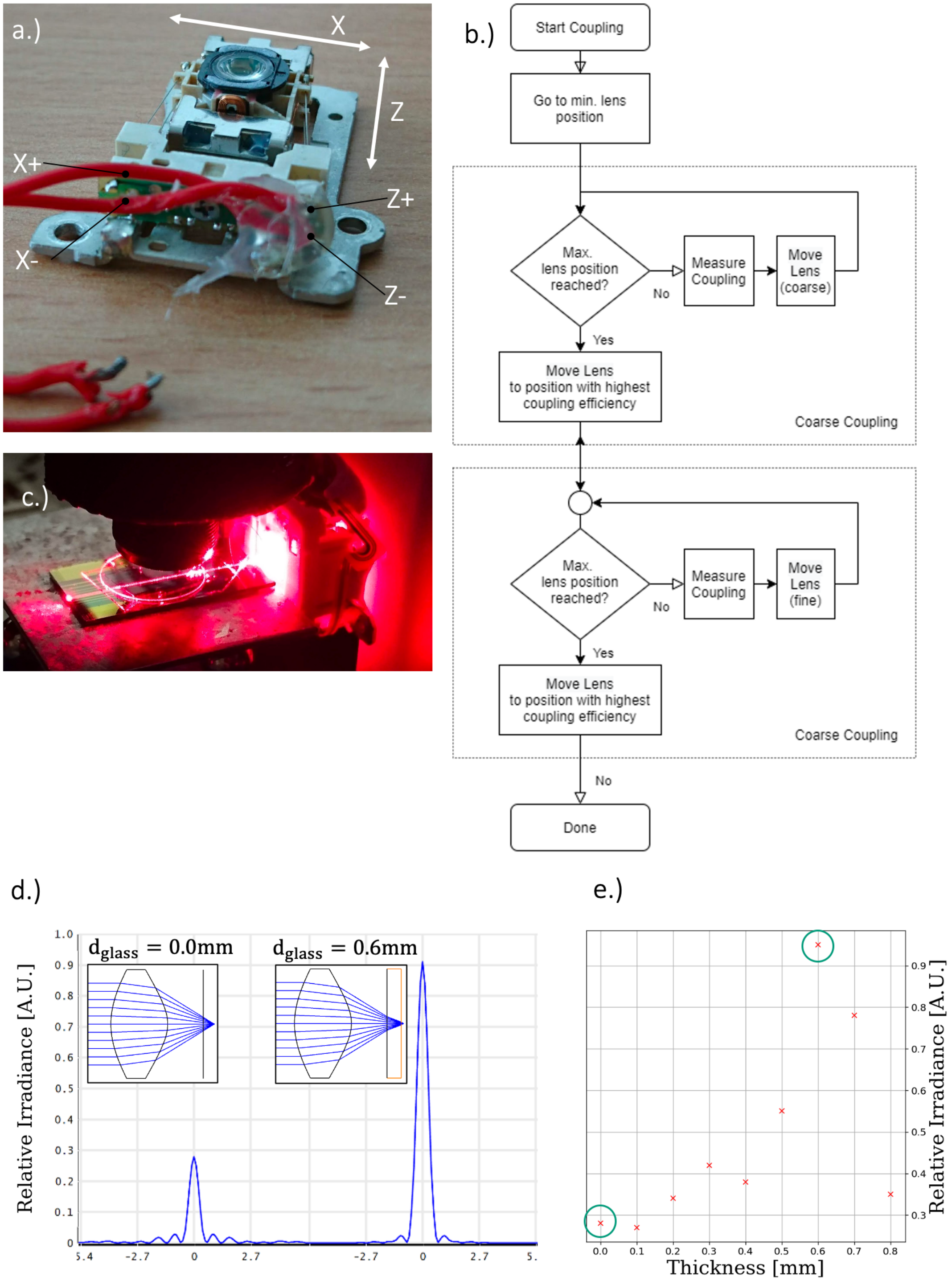


**Figure S5. OPU as accurate coupling device -** The wiring of the OPU in a.) is straightforward. Additionally, the diffraction grating in the back of the lens has to be removed to increase coupling efficiency. A basic coupling algorithm b) increases the signal inside the waveguide c), d) a simulation of adding glass plates between the WG chip and the OPU shows the focussing spot quality. The glass slide (ie.g coverslip) compensates the missing PDMS layer of the disk. d.) For the ZEMAX model, this means a thickness of roughly $0.6\,mm$

#### S1.4 Build instructions

We give a detailed set of instructions, including the 3D printing and design files on our project-webpage.

#### S1.5 ISO vs Noise on P20

In order to give some more insight into the black box like behaviour of the cellphone camera sensor, we characterize its readnoise and gain behaviour for different gains and exposure times.

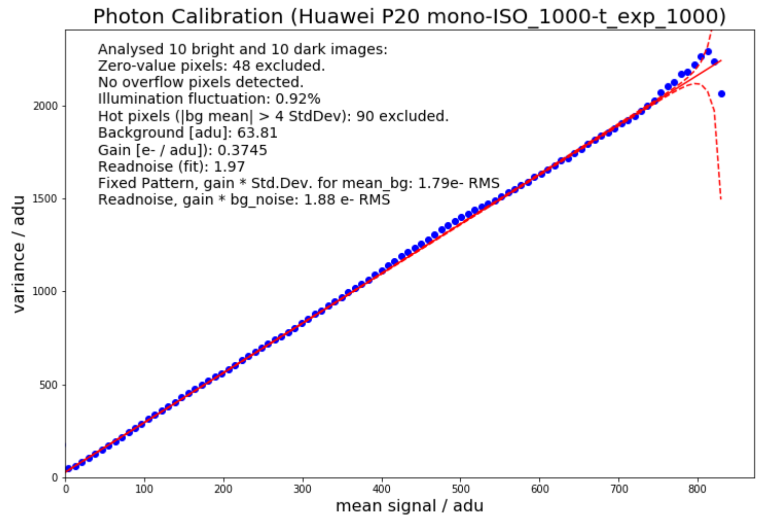


**Figure S6. Flowchart of the autofocus algorithm** - Mean-variance plot generated using a series (ISO = 1000) of unprocessed raw images recorded using the Huawei P20 Pro mono camera (blue points). The camera gain is constant up to an critical intensity of 700 ADU, which should not be exceeded in the experiment. Figure is reproduced from [6]

Therfore, we performed a series of image recording experiments at varying ISO-level and adjusted the exposure time such that the histogram is roughly centred. For each experiment listed in Tab. S1.5, we recorded 10 bright frames of an intentionally defocussed but stationary object which covers a broad dynamic range and dark (i.e. camera fully covered) frames. A customized APP [7] records 12 Bit RAW frames at fixed acquisition parameters.

| **ISO** | **100** | **200** | **500** | **1000** | **3200** | **6400** |
|---|---|---|---|---|---|---|
| $t_{exp}\,[s]$ | 1/60 | 1/125 | 1/250 | 1/1000 | 1/2000 | 1/4000 |
| readnoise | 2.76 | 2.38 | 2.26 | 1.97 | 2.37 | 1.89 |
| Gain $(e^{-}/adu)$ | 2.75 | 1.63 | 0.6752 | 0.3745 | 0.16 | 0.07 |

The mean-variance plot exemplary shown for $ISO = 1000$ and $t_{exp} = 1/1000s$) of the Huawei P20 Pro monochromatic back illuminated CMOS sensor (Sony IMX 600) represents a common setting for fluctuation-based superresolution microscopy. The variance increases linearly with the recorded mean intensity. The gain, estimated as the slope of the curve in Fig. S6 shows an amplification (Gain) of $2,7\,ADU/e^-$ at a readnoise-level of only $1.97e^-RMS$. For ISO-settings above $ISO = 2000$, the number of zero-valued pixels grows dramatically. This could be due to a digital amplification.

It has to be mentioned, that the preprocessing in compressed video-streams, including hot-pixel detection, anti-vignetting and denoising leads to the decorrelation of pixel-information. Therefore, we cannot perform a readnoise calibration on this data, further promoting the need to use RAW-data in cellphone imaging.

#### S1.6 Converting any microscope into a WG-based microscope

The setup here presented, can also be without the optical module and the cellphone. The coupling mechanism then serves as a versatile tool which can for example turn any microscope into a TIRF-ready device as shown in Fig. S7.

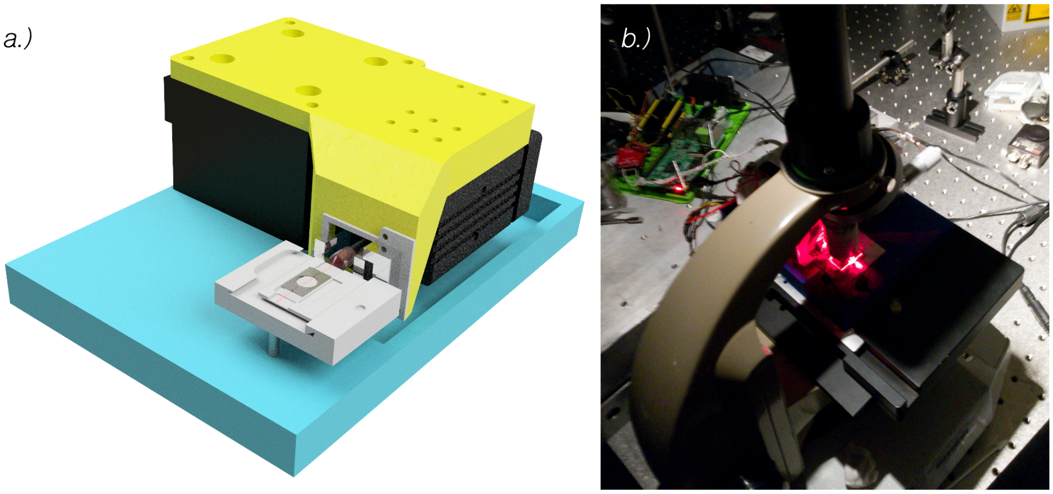


**Figure S7. TIRF conversion of an standard microscope -** Taking only the waveguid coupling unit a) enables a simple conversion from a standard up-right microscope into a TIRF-ready imaging device as shown on an old Wicker tripot b).

##### S1.7 Dual-colour mode for co-localization

In many biological experiments, multiple excitation wavelengths are needed to separate multiple functional groups (e.g. co-localization). Using the dichroimatic mirror from the Bluray drive (KES400A, Sony Japan), we were able to build an adjustable beam-combiner with very little effort S8a). We used the combination of the red (laserlands, see BOM S1.1, China) and green (lily electronics, see BOM S1.1, China) laser in order to excite mCherry expressing pseudo-type virus additionally labelled with Alexa Fluor 647 labelled anti-human antibodies. A kinematically hold folding mirror S8 a)/c) allows placement of the red laser beam inside the back-focal plane of the optical pick-up unit S8a). Therefore we placed a mirror (10 $mm$, Comar optics, UK) glued to a 3D-printed baseplate equipped with three ball-magnets on ferromagnetic screws (M3, Würth, DIN912). This mechanism allows the mirror to be oriented in all directions at high precision and little drift S8c/d).

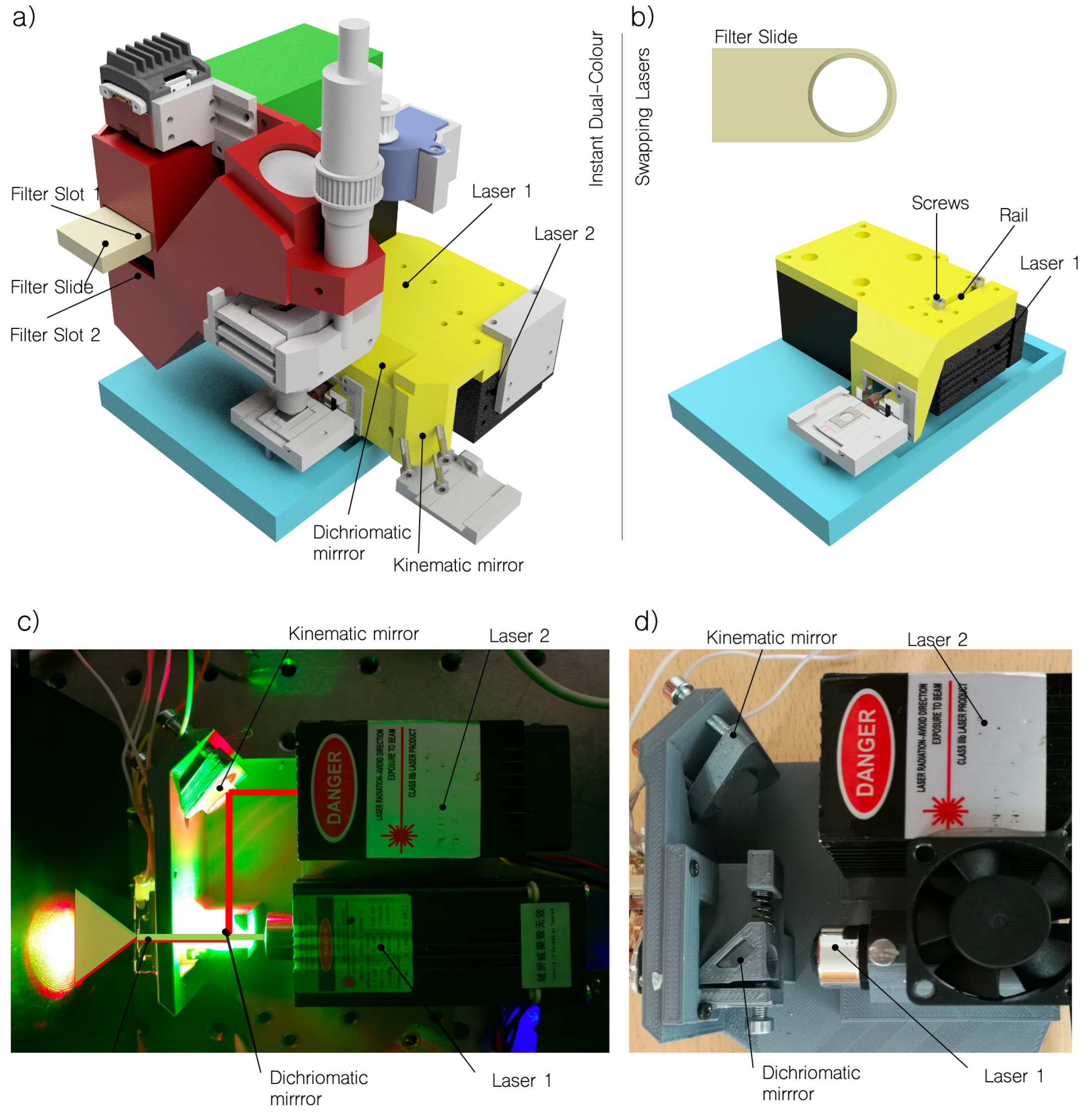


**Figure S8. Dual-colour setup** - a) A dichroimatic mirror from the KES400A represents a low-cost way to create a beam combiner for red ($\lambda = 635\,nm$) and green ($\lambda = 532\,nm$) lasers. A kinematic mirror mount and a stir-able mechanism for the dichroimatic allows an easy aligning procedure to optimize the focusing quality for both colours. A filter slide allows manually inserting different emission filters. b) An easier-to-build dual colour setup provides a 3D-printed rail mechanism which allows mounting a laser by sliding it in and out. c) Aligning the two laser lines is done by observing the image of the laser spot after the OPU while varying the position of the kinematic and dichroic mirror in a). d) Different lasers can be used combined with a different dichroimatic mirror.

The focus for each wavelength slightly differs which can conveniently be compensated by adjusting the OPU's lens position. During the co-localization of the virus particles, we used 3 different wavelength, which resulted in sequentially replacing the lasers in the single-colour setup. This was easily achieved by releasing a single screw and sliding in and out the different laser modules S8b). All laser modules can be controlled using the Android remote control. Filters were replaced by sliding it in and out as well using a filter slide (Fig. S8b). The different configurations can be derived from Tab. S1.7.

| **Fluorophor** | $\lambda_c$ **Laser** | **Power** ($mW$) | **Property** | **Source Laser** | **Filter** |
|---|---|---|---|---|---|
| mCherry | 532 $nm$ | 200 $mW$ | 12V 532mm 200mW green laser dot module fan cooling TTL 0-30kHz long time working | lilly electronics | AHF BrightLine HC 582/75 |
| mCherry | 532 $nm$ | 150 $nW$ | "Grün Laser Modul 532 nm 150 mW Analog" | Pro Laser Systems | AHF BrightLine HC 582/75 |
| GFP | 445 $nm$ | 500 $nW$ | "Blau 445 nm Laser Modul 500 mW Analog" | Pro Laser Systems | Omega Filters 535AF45 |
| GFP | 488 $nm$ | 50 $nW$ | "Blau Laser Modul 488 nm 50 mW" | Pro Laser Systems | Omega Filters 535AF45 |
| Alexa Fluor 647 | 635/637 $nm$ | 300 $nW$ | "Laser diode module TTL stage lighting dj show 12VDC" | Laserlands | AHF HQ 675/55 |

The dual-colour setup in the current version is in an experimental stage. The lack of mechanical stability when inserting different emission filters very often results in a slightly shifted FOV which can be compensated in post-processing (i.e. aligning co-localization of lenti virus and VLP samples). Chromatic aberration of the low-cost objective lenses and the mechanical stress when sliding in/out the filter adds additional defocus and can be compensated with carefully refocussing the sample.

#### S1.8 Effective Pixelsizes

| **Lens** | | $100\times$, 1.25 $NA$ **Oil** | $60\times$, 0.85 $NA$ | $40\times$, 0.65$NA$ |
|---|---|---|---|---|
| Abbe limit: | | 260 $nm$ | 382 $nm$ | 500 $nm$ |
| Imaging Method | RAW (*FreedCam*) | 49 $nm$ | 75 $nm$ | 115 $nm$ |
| | MP4 (2k, *FreedCam*) | 67 $nm$ | 100 $nm$ | 157 $nm$ |
| | MP4 (*STORMImager*) | 131 $nm$ | 205 $nm$ | 320 $nm$ |

### Bibliography


[1] Nils Gustafsson et al. "Fast live-cell conventional fluorophore nanoscopy with ImageJ through super-resolution radial fluctuations". In: *Nature Communications* (2016). ISSN: 20411723.

[2] Christian A. Rothenbach and Mool C. Gupta. "High resolution, low cost laser lithography using a Blu-ray optical head assembly". In: *Optics and Lasers in Engineering* 50.6 (2012), pp. 900–904. ISSN: 01438166. URL: `http://dx.doi.org/10.1016/j.optlaseng.2011.12.004`.

[3] Reuven Rasooly et al. "Improving the Sensitivity and Functionality of Mobile Webcam-Based Fluorescence Detectors for Point-of-Care Diagnostics in Global Health". In: *Diagnostics* 6.2 (2016), p. 19. ISSN: 2075-4418. URL: `http://www.mdpi.com/2075-4418/6/2/19`.

[4] Edwin En Te Hwu and Anja Boisen. *Hacking CD/DVD/Blu-ray for Biosensing*. 2018.

[5] Itseez. *OpenCV 3.0*. 2016. URL: `http://opencv.org/` (visited on 04/27/2016).

[6] Benedict Diederich et al. "UC2 – A Versatile and Customizable low-cost 3D-printed Optical Open-Standard for microscopic imaging". In: *bioRxiv* (Jan. 2020), p. 2020.03.02.973073. URL: `http://biorxiv.org/content/early/2020/03/03/2020.03.02.973073.abstract`.

[7] Benedict Diederich and Ingo Fuchs. *Github: cellSTORM Android APP*. 2018. URL: `https://github.com/bionanoimaging/cellSTORM-ANDROID`.
